# Diversification mitigates pesticide but not microplastic effects on bees without compromising rapeseed yield in China

**DOI:** 10.1101/2024.09.28.615554

**Authors:** Wei Lin, Xueqing He, Jilan Hu, Marcel Balle, Kevin F.A. Darras, Siyuan Jing, Christoph Scherber, Manuel Toledo-Hernández, Yan Yan, ChengCheng Zhang, Siyan Zeng, Thomas Cherico Wanger

## Abstract

Humanity depends on agriculture for food, fiber and energy provisioning, but input-intensive agricultural production is impacting ecosystem services such as pollination. Pollution effects from neonicotinoid insecticides on pollinators receive much attention, but nothing is known on the synergistic effects with emerging plastic contaminants and the mitigation potential of agricultural diversification. Here, we conduct the first large-scale and full-factorial mesocosm study to understand two-generation effects of diversified floral resources (diversification treatment), neonicotinoid and microplastic pollution (pollution treatments) on *Osmia cornifrons* bees in 72 mesocosms. In our three-year experiment, we found that diversification can mitigate individual neonicotinoid effects. We did not find any individual or synergistic effects of microplastic on reproductive performance of solitary bees. None of our treatments affected rapeseed yield. Our results confirm the benefits of diversified flower resources to mitigate pesticide effects on bees in China and suggest that microplastics have no acute individual or interaction toxicity in semi-natural environment at realistic exposure levels. Diversified flower resources in Chinese agricultural landscapes to mitigate pesticide pollution effects on pollinators is an important policy argument for pollinator protection with downstream implications for food security.

## Introduction

Global population increase poses the great challenge upon humanity to produce enough food while minimizing its impacts on the environment under climate change ^1,2^. Agroecology and the underlying approach of agricultural diversification, is one of the promising solutions to tackle this challenge and to achieve sustainable food production^3^. Agricultural diversification includes various practices all focusing on adding functional diversity to enhance ecosystem services within agricultural landscape while at least maintaining yields^4,5^. For example, agroforestry and semi-natural habitats can maintain plant and animal diversity in fields and within agricultural landscapes^6–8^. Moreover, diversifying farming systems by adding non-crop components or around crop field promotes insect diversity and economic benefits especially pollinators^9^. Most angiosperms require animal-vectored pollination^10^, hence threats such as fragmentation of (semi-) natural habitats, overuse of pesticides and environmental pollution to pollination services may compromise food security and ecosystem resilience^11,12^. Farm and landscape level diversification benefits on pollination are well established and have been synthesized in general and for specific crops^6,13^. By offering complementary floral resources over bees’ foraging and extended flowering periods, diverse landscape provides a large variety of nutritional pollen or nectar for bee reproduction^14^. Enhancing flower resources has been shown to mitigate the effects of neonicotinoid pollution on solitary bees^15^. Flower strips in southern Sweden could increase pollinator abundances across agricultural landscapes^16^. The increased availability of flowering plant species can effectively mitigate the negative effects of neonicotinoid insecticides, while double brood cell production compared to crop monocultures^17^. However, little is known about diversification to mitigate the effects of emerging pollutants such as plastic individually and synergistically with neonicotinoids^18^.

Plastic pollution has been increasingly recognized as a major issue in agricultural systems^19^ that is likely going to increase in the future. Depending on the type, size, and concentration, plastic can affect soil properties and plant growth^20,21^ as well as plant microbial communities^22^. Plastic accumulation and retention may be modified by so called ‘NMP islands’ at the farm and landscape scale^18^. These NMP islands are plastic concentration hotspots e.g., at hedge rows or semi-natural habitats that also mediate pollinator diversity and abundance by providing floral resources and nesting opportunities. The high co-occurrence of pesticides and plastics in agricultural systems raises the question of synergistic effects of the two pollutant types. Synergistic effects are known, for instance, in combination treatments of tetracycline and MP that reduce the survival rates in honeybees compared to individual treatments^22^. Plastics can adsorb pesticides^23^, which may lead to overall higher substance concentrations. However, the importance of synergistic effects of highly toxic substances such as neonicotinoids and plastics on pollinators is currently unknown, and mesocosm and field studies are urgently needed.

Here, we tested how diversified flower resources mediate the effects of neonicotinoid pesticides and microplastic on bee reproductive traits and rapeseed traits including yield in a semi-field experiment with 72 mesocosms. We showed that diversification can mitigate individual neonicotinoid effects in Chinese agricultural landscapes. We did not find any individual or synergistic effects of microplastic on reproductive performance of solitary bees and none of our treatments affected rapeseed yield.

## Results

We found that flower resources could mitigate neonicotinoid effects on bee brood cells and cocoons (Fig. 1A, 1B, Tab. S3, Tab. S7) and that microplastic did not affect bee traits (Fig. 1C, Tab. S3, Tab. S7). In mesocosms without diversified flower resources, brood cell numbers (Fig. 1A, P < 0.001; Table S12) and cocoon numbers (Fig. 1B, P < 0.01; 1C, Tab. S12) of *Osmia cornifrons* were significantly reduced by neonicotinoids. The proportion of daughters in the F_1_ generation was not affected by any treatment (Fig. 1C, Tab. S3, Tab. S7). Mechanistically, flower resource diversity increased brood cell and cocoon counts measured as Shannon Diversity (cells number: P<0.001, cocoons number: P<0.01) and high flower species richness (cells number: P<0.001, cocoons number: P<0.01; Fig. 2, Tab. S11).

**Figure. 1.**
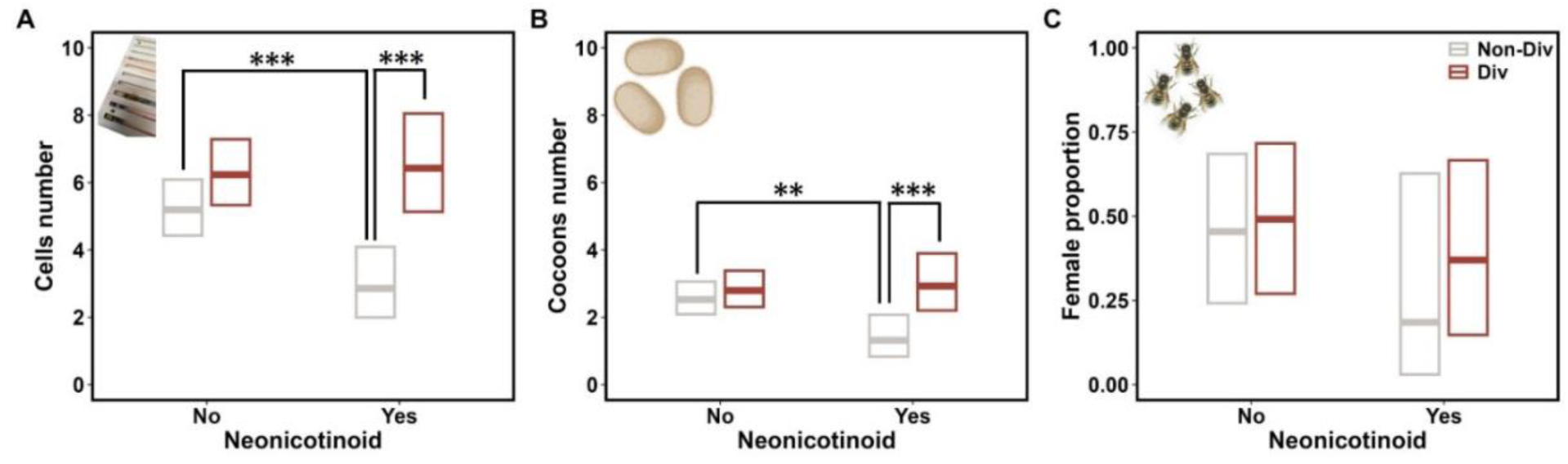
Effects of neonicotinoids and diversification on (A) bee brood cell number, (B) cocoons number and (C) female proportion in the F_1_ generation. Predictions and 95% confidence intervals come from generalized linear mixed models. Asterisks denote significant differences between groups with Type-III analysis of variance (* P≤0.05; ** P≤0.01; *** P≤0.001).

**Figure. 2.**
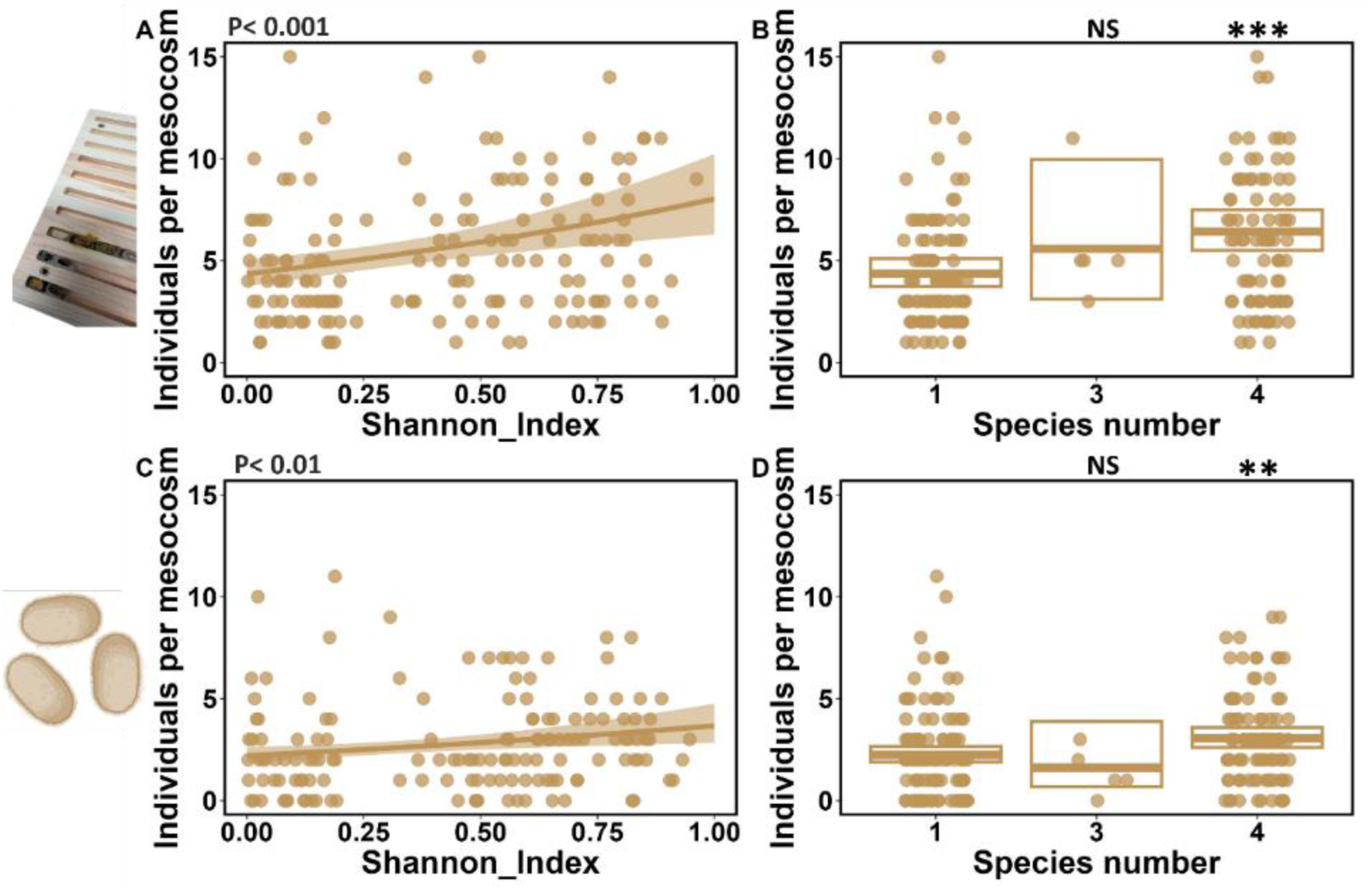
Bee brood cell and cocoon counts per mesocosm increase with flower resource diversity as measured by (A, C) shannon index and (B, D) species number per mesocosm. Predictions and 95% confidence intervals come from generalized linear mixed models (NS = not significant; * = P≤0.05; ** = P≤0.01; *** = P≤0.001).

We did not find effects of pollutant treatments on rapeseed flower number, as diversified flower resources intuitively decreased rapeseed flower number (Fig. S3. A, Tab. S5, Tab. S6). We did not find any effect of the pollutant and diversification treatment on rapeseed yields (Fig. 3, Tab. S4, Tab. S8). Neonicotinoids had no effect on rapeseed plant dry weight and rapeseed pod number, but the microplastic treatment increase rapeseed plant dry weight when flower resources were enhanced (Fig. 4, Fig.S5, Tab. S4, Tab. S13). We did not find any effect on pod dry weight and pod *Alternaria* disease incidence (Fig. S5. B, S6, Tab. S4, Tab. S8).

**Figure. 3.**
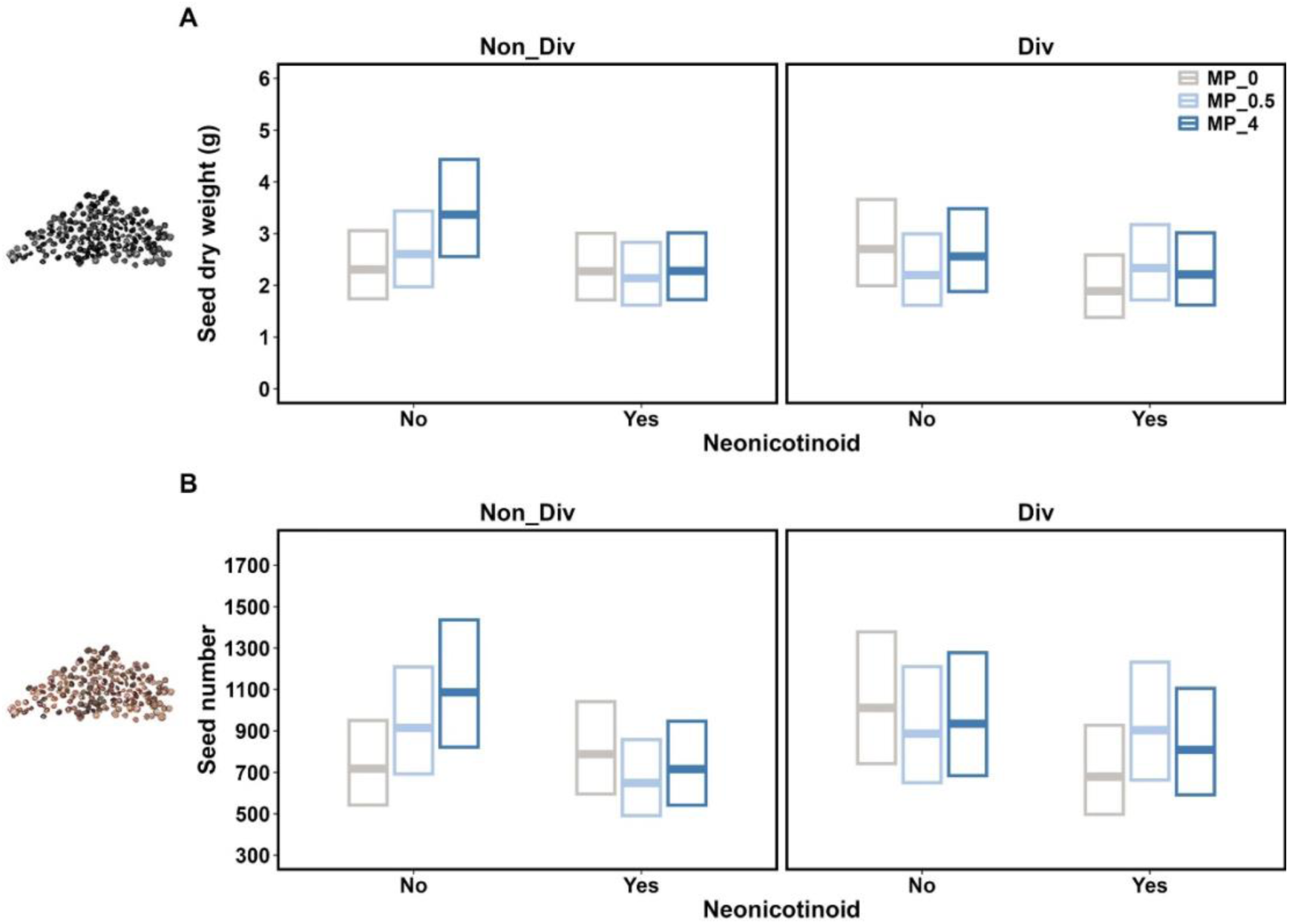
Effects of diversification, neonicotinoids and microplastics on (A) Seed number, (B) Seed dry weight. Predictions and 95% confidence intervals come from generalized linear mixed models. Asterisks denote significant differences between groups with Type-III analysis of variance (* P≤0.05; ** P≤0.01; *** P≤0.001). Treatment abbreviations: MP_0= soil containing 0g/kg microplastic; MP_0.5 = soil containing 0.5g/kg microplastic; MP_4.0 = soil containing 4.0g/kg microplastic.

**Figure 4.**
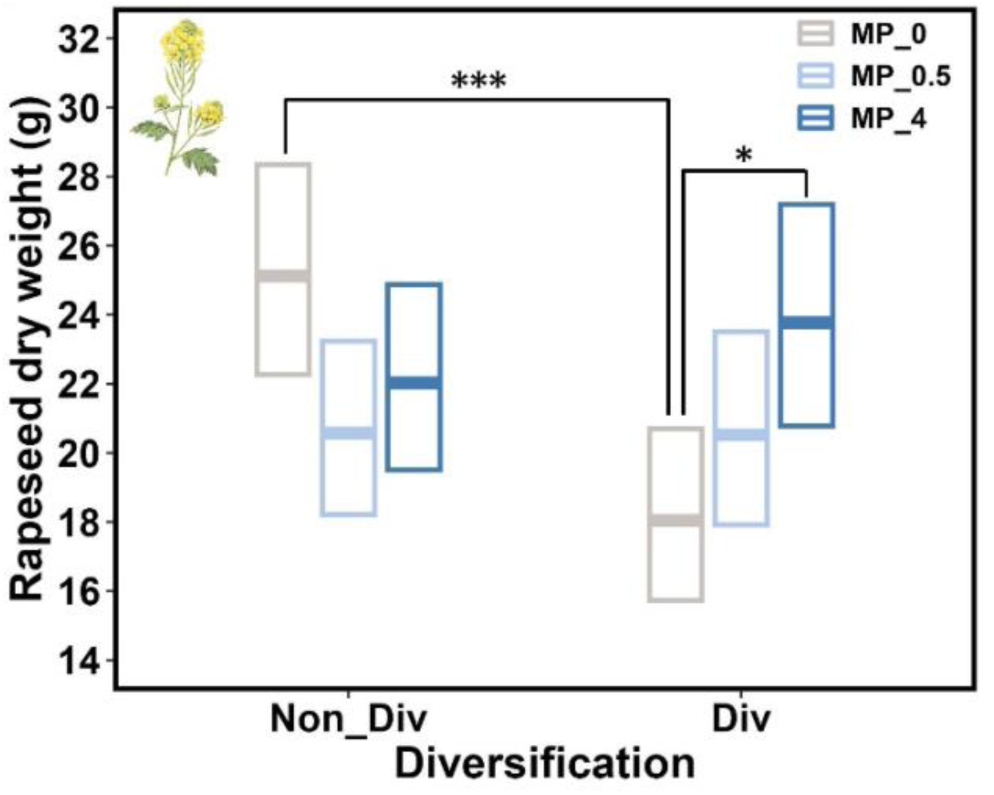
Effects of microplastic and diversification on rapeseed plant dry weight. Predictions and 95% confidence intervals come from generalized linear mixed models. Asterisks denote significant differences between groups with Type-III analysis of variance (* P≤0.05; ** P≤0.01; *** P≤0.001). Treatment abbreviations: MP_0= soil containing 0g/kg microplastic; MP_0.5 = soil containing 0.5g/kg microplastic; MP_4.0 = soil containing 4.0g/kg microplastic.

## Discussion

### Diversification mitigates pesticides effect on bees

Pollinators are declining from pressures resulting from intensive agriculture such as pesticide overuse and plastic pollution^24,25^, the individual and synergistic effects as well as the mitigation potential of diversified flower resources remain poorly understood^18^. We showed that flower resource diversification can mitigate effects of neonicotinoids on pollinators in China. The number of nests and cocoons grew with all metrics of flower species diversity, a diverse pool of complementary resources for food and nesting^17^. The solitary blue orchard bee (*Osmia cornifrons*) made fewer, longer foraging trips and often misidentified nests with limited food resources, while neonicotinoid exposure reduced their foraging activity^15,26^. The observed effects of neonicotinoid on pollinators may be mitigated in mesocosms with diversified flower resources, because extra and attractive floral resource potentially decrease bee contact with pollutants^17^. In the field, flower strips have for instance been shown to attract and support bees in intensively used agricultural landscapes and limit exposure to pollen and nectar containing neonicotinoid^27^.

Our results suggested that microplastics do not have significant effects on bee colonies either individually or in synergy with neonicotinoids. Given the increasing exposure of bees to both, pesticides and microplastics in agricultural landscapes, this is a reassuring perspective. However, there is evidence that neonicotinoids can be adsorbed on plastic agricultural film particles^28^, which may increase the exposure of bees to neonicotinoid from pollen or the soil. Other studies show that a combined exposure to pesticides and other pollutant, which is driven by agricultural intensification, contribute to the decline of bee colonies^29^. Despite we did not find individual or synergistic effects of microplastic on the sex ratio of the F_1_ generation, future studies should investigate combined plastic-neonicotinoid effects on bee sex ratio. A decline in the proportion of females in honeybees affects the subsequent development of the population and leads to colony decline^30^. Overall, our work is the first to investigate microplastic and neonicotinoid effects on bees under semi-field conditions in China. More research is urgently needed to investigate these effects in different parts of the world with different plastic types, sizes, and concentrations^18^.

### Diversification and pollution treatments are not affecting rape seed yield

Our results confirm findings from other studies that diversified flower resources do not affect rapeseed yield^31^, a phenomenon that also extends to non-crop diversification in other crops^32^, across regions^33^, and cropping systems in general^34,35^. We also did not find any effects of neonicotinoids and flower resource diversification individually or in combination on other rapeseed traits. The diversified flower resources maintained rapeseed yield through reducing negative effects of pesticides on pollination service. As systemic insecticides, neonicotinoids are distributed equally across the plant and defending against sucking and chewing insect pests. The lack of effects of neonicotinoids on rapeseed traits suggests that pest pressure in our research site may not be so high that neonicotinoids use is resulting the anticipated benefits. As neonicotinoids cause pollinator population decline, the use of these insecticides as a standard practice in our study region should be carefully considered. We also found that high microplastic concentrations increased rapeseed plant dry weight. This result is in line with other field studies^38^, but such effects are highly species and location dependent^20^. Future research should concentrate on the response of various crop species to realistic exposure concentrations of microplastics in agricultural system.

### Conclusions

While neonicotinoid are commonly known threats to pollinators^39,40^ and microplastic pollution is a rapidly emerging threat to the environment^41^, research on interaction toxicity between these pollutants and particularly in the field is still in its infancy. Our study makes an important first step to address this research gap with ramifications for food security. Future research should focus on the effects of different plastic types of various sizes and their interactions^42^, to then understand the underlying mechanisms between the plastic transfer in the soil-plant-pollinator system. New methods^32^ need to be developed and applied to facilitate analyses of how local conditions in climate, soil type, and crop as well as pollinator species affect the plastic transfer. Lastly, as diversification is increasingly proposed as a solution towards the global food system transformation in national and international policy ^43, 2^, our work represents a first step to support policy advisory on pollutant mitigation through effective diversification strategies such as flower strips.

### Methods Study sites

The experiment was conducted in Fuyang district, Hangzhou City (30.13 °N, 119.95 °E), the northwest of Zhejiang Province. Fuyang district has a subtropical monsoon climate with an average annual precipitation of 1426−1800 mm, average temperature of 14-23 ℃, frost-free period of 230-260 days, and 1791 sunshine hours. Forest cover amounts to 63 % and the topography is hilly. The main crops in the area include rice, rapeseed, vegetables and tea. The local soil types are mainly red and yellow earths, paddy soil (Ao 13-3bc category in FAO/UNESCO Soil Category). The study field has always been planted with rice, fallowed during winter or rotated with rapeseed. General management consisted of a basic fertilizer application before the planting and one topdressing, and two pesticide applications. Neonicotinoid was generally used at our experimental concentrations.

### Flower resource, Neonicotinoid and Microplastic Treatments

The experiment was planned as a full factorial design with each microplastic treatment being combined with and without a neonicotinoid treatment, and random treatment allocation in the field (Fig. S2). We used a mesocosm experiment of 72 cages covered with Nylon fine mesh (hereafter mesocosms) of 2m × 4m × 2m from October 2021 until May 2022 for the experiment. Thirty pots (60cm × 24cm × 23cm; PVC) per cage were prepared with soil collected from the site, where the experiment was conducted. We used soil from the site because it was - after several attempts - logistically not feasible to order the required quantity of restored (i.e., completely unpolluted) soil for the experiment. The soil and water pollution chemical background check (conducted by ICAS Testing Center, Shanghai, China) showed no pollution of neonicotinoids and microplastics (Extended data Fig. 1, Extended data Fig. 2). The soil was then mixed and equally distributed to all pots. Our diversification treatment to diversified flower resources in half of the mesocosms (36 cages) contained 50% (15 pots) autumn rapeseed (variety ‘Zheda630’, commonly cultivated in the local area) and 50% (15 pots) flowering plants seed mixture. The remaining mesocosms (36 cages) were planted with 100% (30 pots) rapeseed as a control to the diversification treatment (Fig. 5). The diversification treatment contained six plant species, selected to flower at the same time as the autumn rapeseed (March/April) and to be attractive to our target pollinator, *Osmia cornifrons*: Chinese Milk vetch *(Astragalus sinicus)),* Chinese violet cress *(Orychophragmus violaceus),* Cornflower *(Centaurea cyanus),* Corn poppy *(Papaver rhoeas),* Zinnia *(Zinnia elegans)* and Broad bean *(Vicia faba)*. As a substitutive design, we planted three rapeseed plants in one pot, and the six flowering plants were mixed and sown in other pots. Our diversification treatment successfully increased plant richness, flower diversity (Fig. 2), and – due to the nature of the flower strips – did not increase flower numbers (Fig. S3 B, C).

**Figure. 5.**
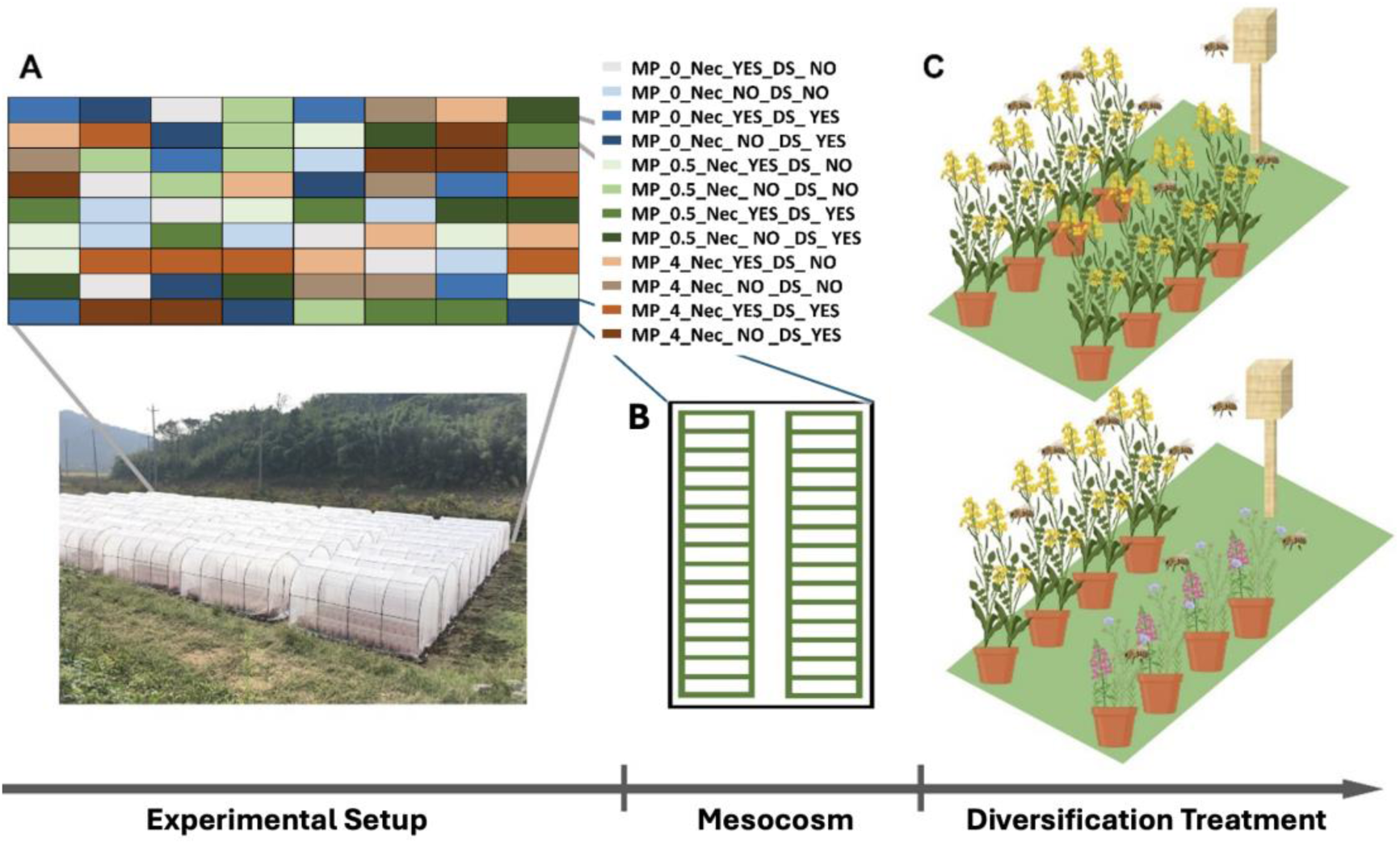
Experimental Setup including treatment distribution and schematic of the diversification treatment. Treatment abbreviations: MP_0.5 = soil containing 0.5g/kg microplastic; MP_4.0 = soil containing 4.0g/kg microplastic; Nec_YES = rapeseed seeds treated with neonicotinoid; Nec_NO = rapeseed seeds not treated with neonicotinoid; DS_YES = diversification treatment, flowering plants planted in cage; DS_NO = no diversification treatment and no flowering plants planted in cage.

We planted neonicotinoid-treated (*Clothianidin*; 5g/kg) autumn rapeseed seeds in half of the mesocosms (36 cages) at normal application rates for the study area and recommended application dosage. Our two micro-plastic treatments followed realistic exposure concentration (0.5 g/kg) and a concentration where reduced plant growth was observed^44^ (4 g/kg). We weighed each pot and measured soil humidity, to then calculate the dry weight of soil in each pot and add the micro-plastic treatments concentrations of 0.5 and 4 g/kg. The micro-plastic was spread evenly on the soil surface in each pot, and then mixed into the soil with a small shovel. We utilized polyethylene (PE) microplastics in the form of spherical particles ranging from 380 to 1000 µm. As this size is too big to be absorbed into plants and carried from the roots to the pollen and nectar, the effects on plants and bees should be solely due to changed plant growth and the number of floral resources produced. We specifically used PE instead of PVC (polyvinyl chloride) because PE is widely found in agricultural systems^45^.

We followed the common rapeseed management practice in the study area for all mesocosms. We applied a base chemical fertilizer before putting the soil into the pots and a topdressing when the rapeseed plants had five leaves. We applied the herbicide Acetochlor (active ingredient: acetochlor) after sowing the rapeseed, and one fungicide Carbendazim (active ingredient: methyl benzimidazole carbamate) during the experiment to control fungal infections. There was no artificial irrigation and the whole experiment relied on rainfall.

### Bee reproductive success

The solitary bee species *Osmia cornifrons*, a Japanese/Asian relative of the European *Osmia cornuta,* is commonly used for agrochemical product testing^46^. The cocoons of *O. cornifrons* were bought from Yantai Bifeng breeding sales center and stored in 4 °C before the blooming of rapeseed. After keeping cocoons at 25 °C for 2-3 days, bees hatched and were introduced into mesocosms on 19 March 2022. In each mesocosm, 10 females and 8 male bees foraged, mated, and nested without human interference.

We installed custom-made wooden hives in every mesocosm before releasing the bees (Fig. S1. B). A hive consisted of 5 nesting boards with 10 nesting cavities (8 mm diameter) and a glass instead of plastic cover to avoid PE contamination of the experiment. On the side of the mesocosm where the hive was installed, we provided sufficient mud and water for bees to build nests and drink. As the lifespan of adult *O. cornifrons* is around 1 month, we collected all nest boards and stored them in a dry indoor environment on 19 April 2022 after all nesting behavior had ceased. Of all the 72 hives deployed, we collected a total of 206 wooden nesting boards with an average of 2.8 nesting boards per hive. The bee brood cells on each nesting board were counted, and then the nesting boards were stored at room temperature in the lab to allow the larvae to develop. In December 2022, cocoons were counted and moved from nest boards to glass containers wrapped with gauze and rubber bands (Fig. S2. D) ^17^. In March 2023, we found that all bees had died in the cocoons. We identified the gender of each adult of the F_1_ generation by morphology: larger female bees compared to males have reddish-brown hairs on the abdomen^17^. The sex ratio was calculated as 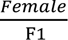

### Flower resources

Flower numbers of all species in each mesocosm were estimated at 3-4 day intervals during the period from March 24^th^ to April 11^th^, 2022, when bees lived in the mesocosms. In pure rapeseed mesocosms, the number of open flowers in 10 randomly selected pots were counted and the average flower number per pot was calculated to estimate the total amount of flowers mesocosm. The same method was applied for diversification cages where the flower number was recorded from 5 rapeseed and 5 diversity pots.

### Plant yield and dry weight

We harvested all plant materials on 19 May 2022 when more than 2/3 of all the rapeseed pods in the field had turned yellow following the general rapeseed harvesting practice^47^. As harvesting and processing all plants was logistically not feasible in the field, given the available time, resources and outside temperature, we chose a random subset of six pots per mesocosm: in the rapeseed mesocosms, 18 rapeseed plants from 6 pots were harvested; in diversification treatment mesocosms, 9 rapeseed plants from 3 pots and all plants from 3 diversity pots were harvested. We marked all plants harvested from the same pot, uprooted them and cleared them from the soil.

To determine rapeseed yield, we counted the total number of rapeseed pods and the number of diseased pods with *Alternaria* disease per plant in the lab. The latter was important to assess how disease incidence was modified through the diversification treatment. For dry weight data, all plant pods were dried in the oven at 70°C (DKM410VC, Zhejiang Creda Scientific Instrument Co., Ltd.) for 48 hours and weighted. To obtain the dry yield, the individual rapeseeds were extracted manually and weighted. We minimized human sampling bias by following a strict sampling protocol. We did not use PE plastic for sampling materials (e.g., sampling bags) to minimize non-treatment related contamination.

### Statistical analysis

All statistical analyses and data visualizations were performed in R^48^ (version 4.3.1). As our experiment was laid out as a randomized complete blocks design, we used generalized linear mixed-effects (GLMM) models and generalized additive mixed model (GAMM) for analysis with Cage ID as a random effect with 72 levels. GLMM models were fitted using Template Model Builder (TMB) implemented in the R package *glmmTMB*^49^ (version 1.1.9). GAMM models fitted with gamm4 based on gamm from package mgcv^50^ were converted to gamViz objects in mgcViz^51^ package with gammV function.

We started off with a design-based analysis, where the three experimental factors were included as fixed effects into the models (including their three-way interaction) as Microplastics (0, 0.5 or 4 g/kg) x Diversification (No or Yes) x Neonicotinoids (No or Yes). Continuous response variables (plant weight, pod dry weight, seed dry weight) were analyzed using a Tweedie distribution^52^ accounting for a range of power-variance relationships in generalized linear models. Briefly, depending on the Tweedie parameter ξ, the Tweedie exponential dispersion model (EDM) allows modeling mean-variance relationship according to the equation

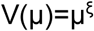

where V is the variance and µ is the mean, with ξ ranging from 0 to 3. Depending on the estimated value of ξ, these models can flexibly account for overdispersion in continuous response variables, ranging from a Normal distribution (ξ=0, i.e. V(µ)=1) to a Gamma distribution (ξ=2, i.e. V(µ)= µ²).

Count data were analyzed using a negative binomial distribution with a quadratic parameterization^53^ according to

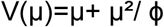

where ϕ is the dispersion parameter (an exponential function of the linear predictor, ϕ =exp(η)).

Proportion data (sex ratio, pod disease incidence) were analyzed using binomial models fit by Penalized Quasi-Likelihood in the glmmTMB library in R^48^ as beta-binomial models in glmmTMB did not converge. Flower number data (rapeseed, diversified flower number) with multiple measurements over time were analyzed using GAMM with a negative binomial distribution in the mgcViz^51^ library in R^48^.

Significance of terms in these models was assessed using type-III analysis of deviance tables for single model objects based on Wald Chi-square tests, where the significance of each term is tested in presence of all others (obeying to the principle of marginality), resulting in order-independent tests, implemented in the Anova() function in the car library in R^54^. After the ANOVA test, pairwise comparisons using Tukey’s HSD adjustment were conducted with emmeans library in R^55^.

After having identified the effects of main experimental treatments, we ran additional models with covariates such as plant diversity to explore the mechanisms of the diversification treatment on pollinator-related variables. We tested the effects of increasing floral resource diversity measured as Shannon diversity and richness of species (based on max flower number of rapeseed and all flower species in mesocosm) on bee-related variables.

## Acknowledgements

This work was funded by a Westlake University Start-up grant (TCW).

## Author contributions

Conceptualization: X.H., T.C.W.; Data collection: W.L, X.H., J.H, M.B., K.F.A.D., S.J., M.T.H., Y.Y., S.Z., T.C.W.; Methodology and analyzes: W.L., X.H., C.S., C.Z., T.C.W.; Visualization: W.L.; Writing – original draft: W.L., X.H., J.H., T.C.W.; Writing – review & editing: all authors; Funding acquisition: T.C.W.; Project administration: T.C.W.; Supervision: T.C.W.

## Supplementary Material

### Figures

**Figure S1.**
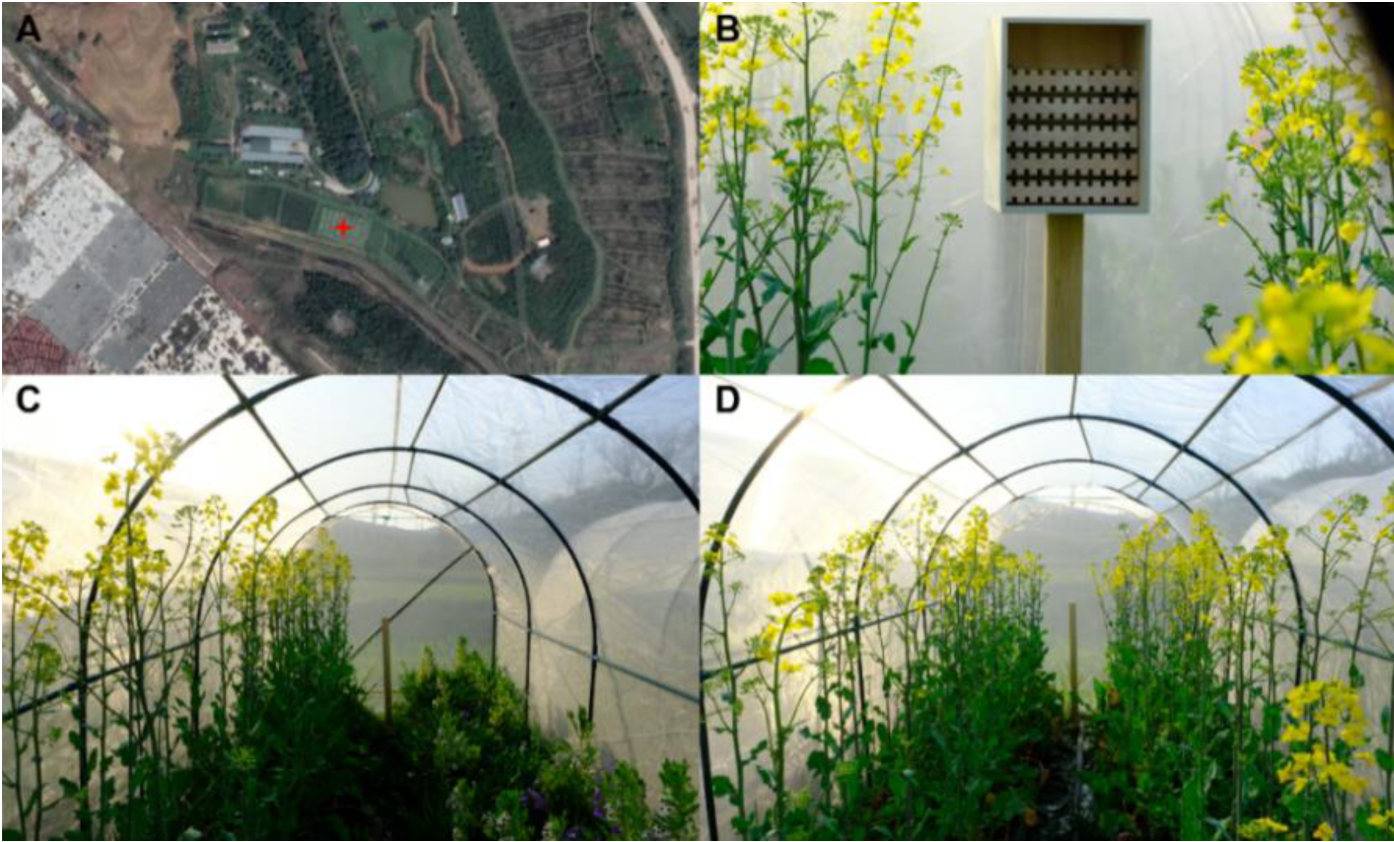
(A) Study site location in Fuyang district, Hangzhou City, Zhejiang Province. (B) Bee nests consisting of 5 nesting boards with 10 nesting cavities in mesocosms. (C) Diversification treatment mesocosms. (D) Non-diversification treatment mesocosms.

**Figure S2.**
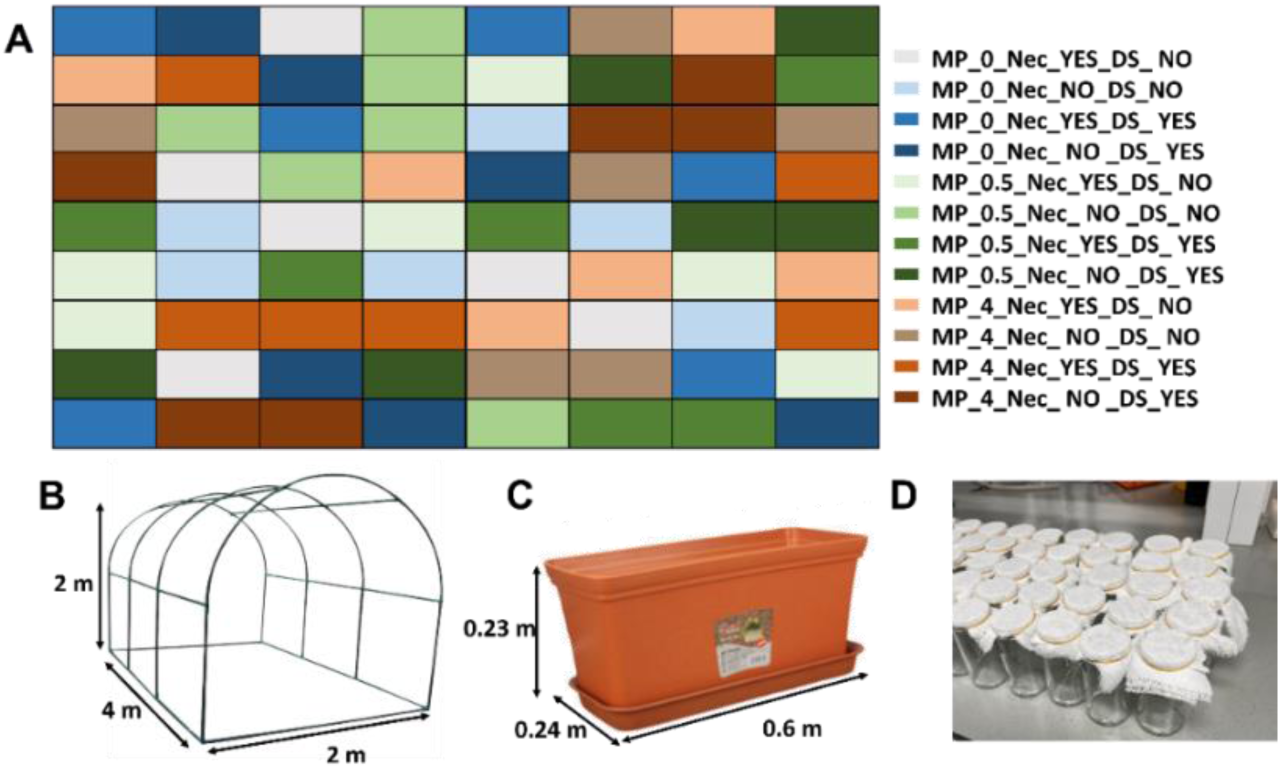
(A) Random treatment layout in the field. (B) Mesocosm size parameters. (C) Pot size parameters. (D) Glass containers with non-plastic cloth covers for storing cocoons. Treatment abbreviations: MP_0.5 = soil containing 0.5g/kg microplastic; MP_4.0 = soil containing 4.0g/kg microplastic; Nec = rapeseed seeds treated with neonicotinoid; MP_0.5_Nec = 0.5g/kg microplastic and rapeseed seeds treated with neonicotinoid; MP_4.0_Nec = 4.0g/kg microplastic and rapeseed seeds treated with neonicotinoid; Control = no microplastic or neonicotinoid application.

**Figure S3.**
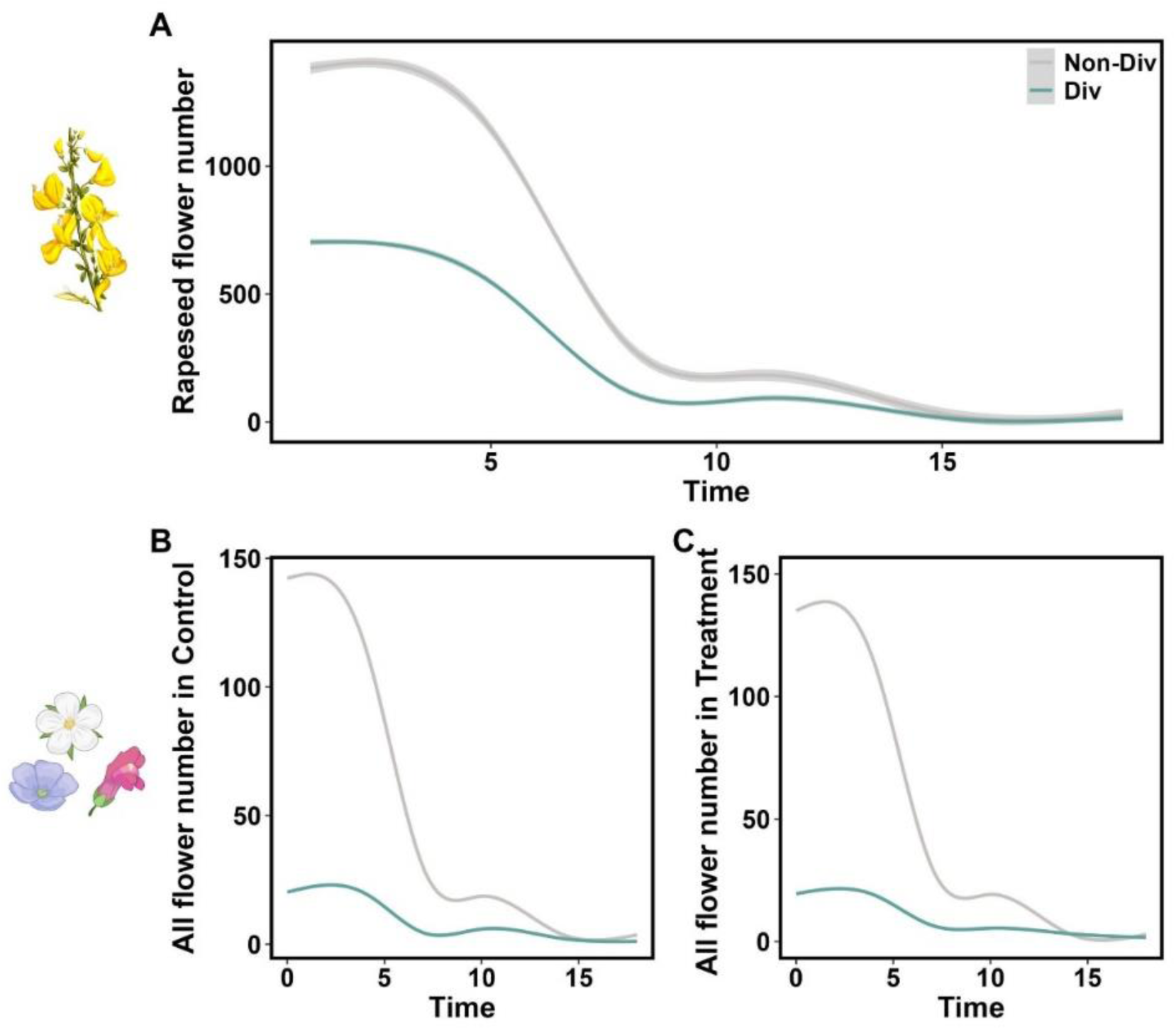
Flower numbers over time. (A) Rapeseed flower number over time in three-way interaction model; (B) All species flower number over time in Control; (C) All species flower numbers over time in all treatments. Predictions and 95% confidence intervals come from generalized additive mixed models.

**Figure. S4.**
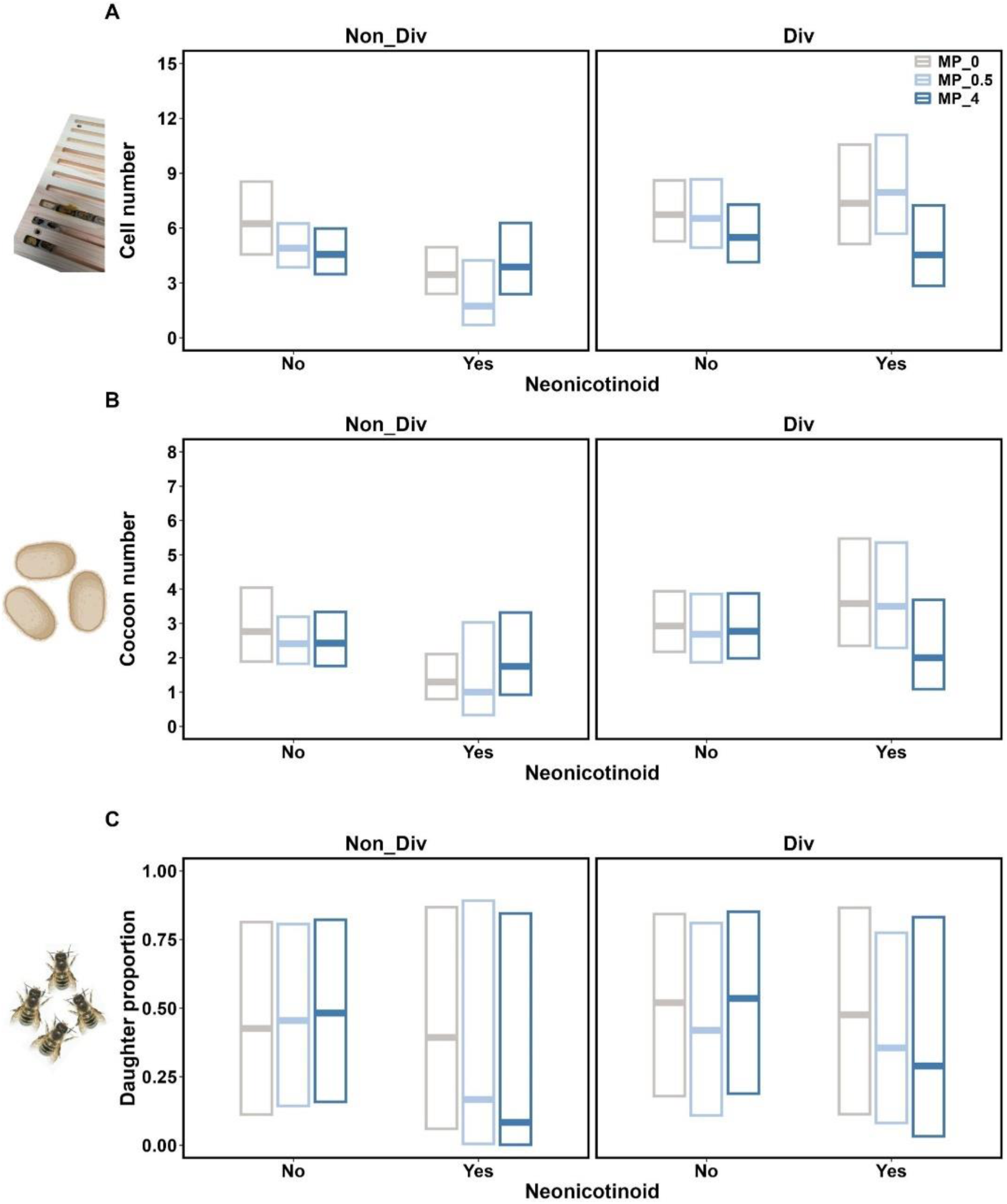
Full factorial treatment effects on bee. (A) bee brood cell number, (B) cocoons number and (C) F_1_ female proportion. Predictions and 95% confidence intervals come from generalized linear mixed models. Treatment abbreviations: MP_0= soil containing 0g/kg microplastic; MP_0.5 = soil containing 0.5g/kg microplastic; MP_4.0 = soil containing 4.0g/kg microplastic.

**Figure. S5.**
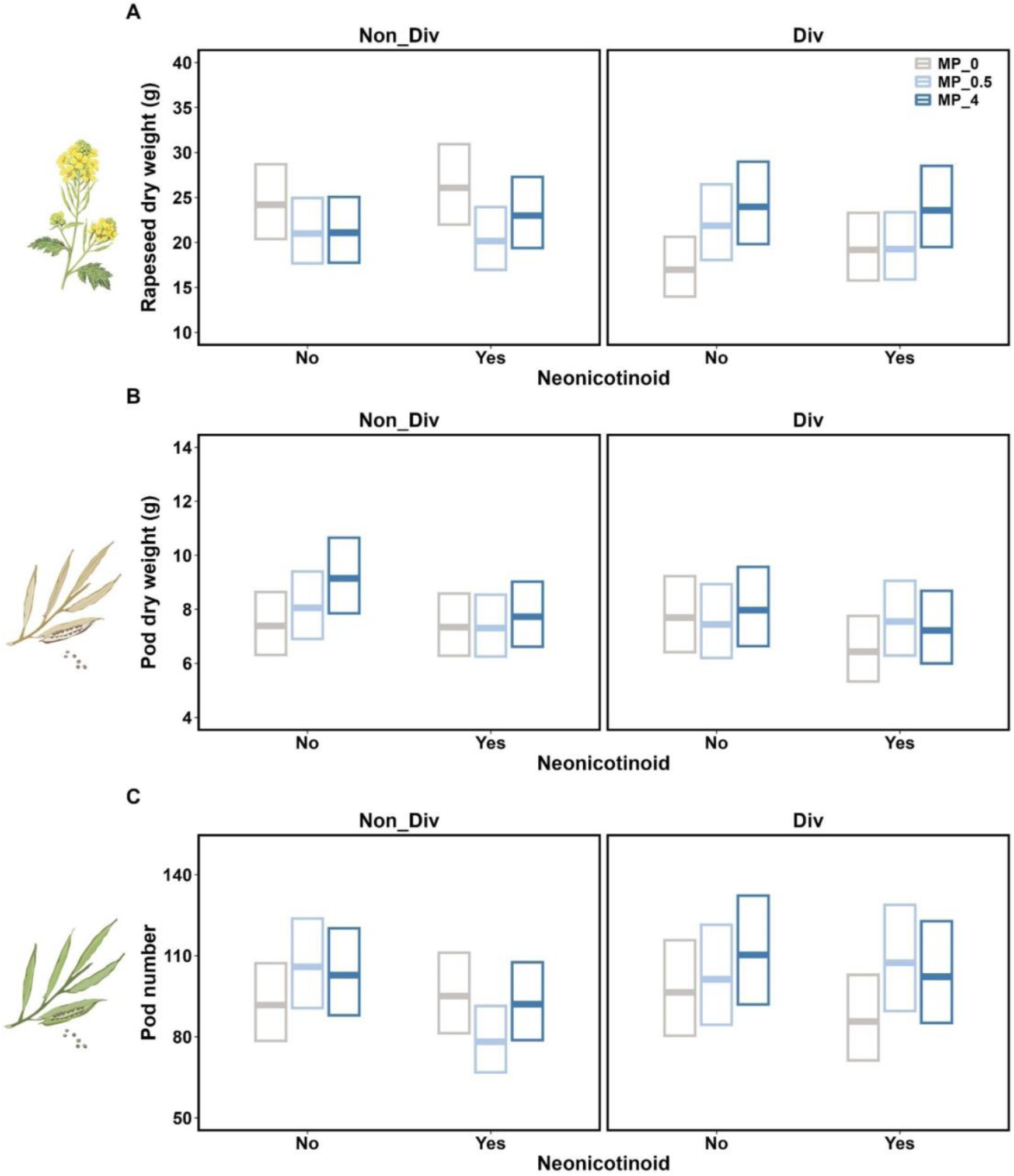
Full factorial treatment effects on rapeseed. (A) rapeseed plant dry weight, (B) pod dry weight. (C) pod number. Predictions and 95% confidence intervals come from generalized linear mixed models. Treatment abbreviations: MP_0= soil containing 0g/kg microplastic; MP_0.5 = soil containing 0.5g/kg microplastic; MP_4.0 = soil containing 4.0g/kg microplastic.

**Figure. S6.**
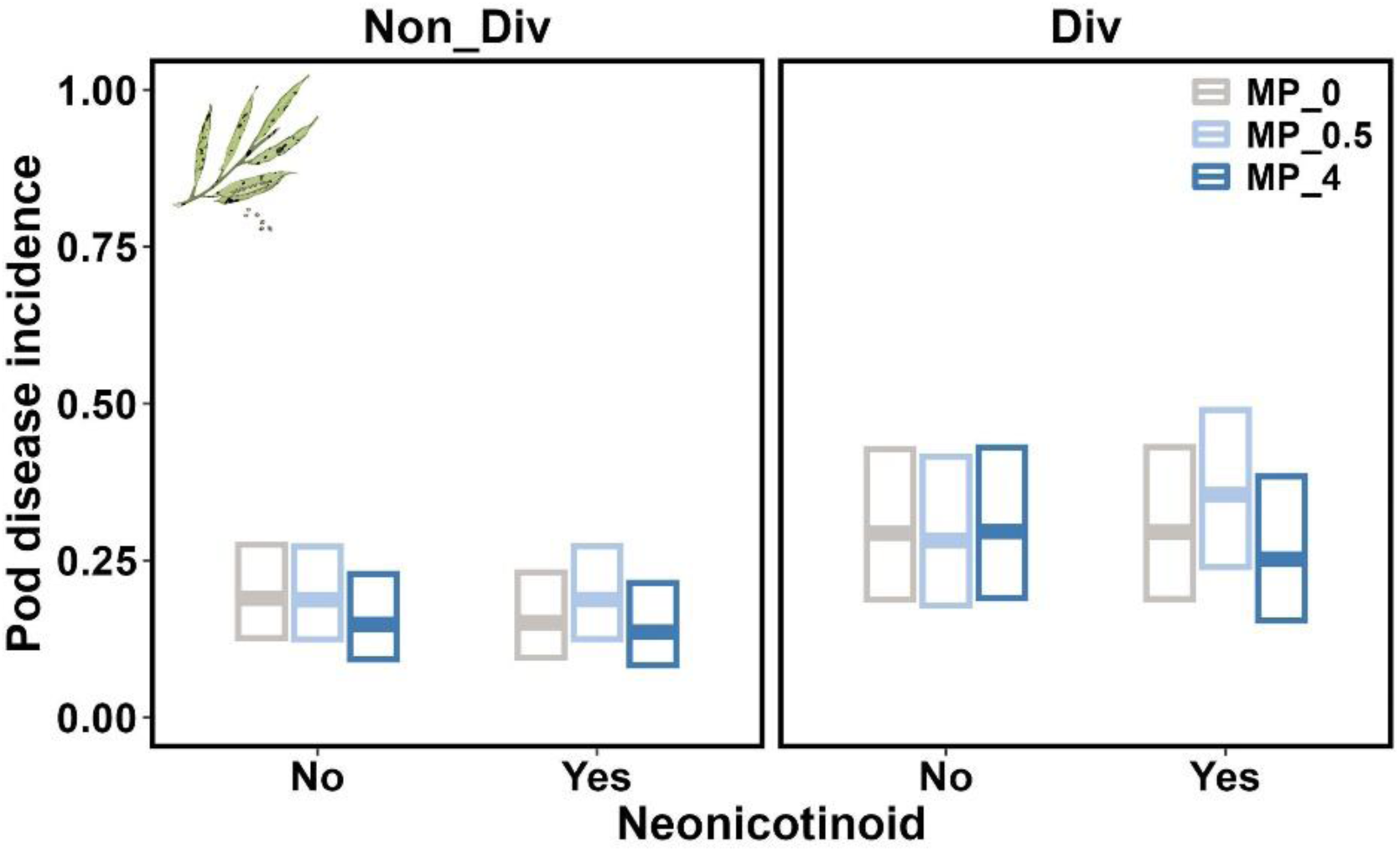
Full factorial treatment effects on pod *Alternaria* disease incidence. Predictions and 95% confidence intervals come from generalized linear mixed models. Treatment abbreviations: MP_0= soil containing 0g/kg microplastic; MP_0.5 = soil containing 0.5g/kg microplastic; MP_4.0 = soil containing 4.0g/kg microplastic.

### Tables

**Table S1.** Properties of the soil used for the experiment in the field site (Extended data Fig. 3). Values in bold are below the indicated detection limit. Soil analysis did not show any neonicotinoid pollution or microplastic fragments (Extended data Fig. 1).

| Soil property | Value | Unit | Soil property | Value | Unit |
| --- | --- | --- | --- | --- | --- |
| pH | 5.7 | - | β-Endosulfan | **<0.09** | mg/kg |
| Total nitrogen | 1.8 | g/kg | Endosulfan sulfate | **<0.07** | mg/kg |
| Total phosphorus | 0.325 | g/kg | α-Lindane | **<0.02** | mg/kg |
| Soil organic matter | 35.2 | g/kg | β-Lindane | **<0.02** | mg/kg |
| Hg | 0.075 | mg/kg | γ-Lindane | **<0.02** | mg/kg |
| Cd | 0.13 | mg/kg | δ-Lindane | **<0.02** | mg/kg |
| As | 5.77 | mg/kg | p,p'-DDE | **<0.02** | mg/kg |
| Pb | 27.6 | mg/kg | o,p'-DDE | **<0.02** | mg/kg |
| Cr | 82.7 | mg/kg | p,p'-DDD | **<0.02** | mg/kg |
| α-Endosulfan | **<0.06** | mg/kg | p,p'-DDT | **<0.02** | mg/kg |

**Table S2.** Water property in the field site (Extended data Fig. 4).Values in bold are below the indicated detection limit. Water analysis did not show any neonicotinoid pollution or microplastic fragments (Extended data Fig. 2).

| Water property | Value | Unit | Water property | Value | Unit |
| --- | --- | --- | --- | --- | --- |
| pH | 7.5 | - | Endosulfan sulfate | **<0.04** | ug/L |
| Total nitrogen | 1.48 | mg/L | α-Lindane | **<0.06** | ug/L |
| Total phosphorus | 0.98 | mg/L | β-Lindane | **<0.04** | ug/L |
| Hg | **<0.04** | ug/L | γ-Lindane | **<0.03** | ug/L |
| Cd | 3.0 | ug/L | δ-Lindane | **<0.06** | ug/L |
| As | 4.1 | ug/L | p,p'-DDE | **<0.04** | ug/L |
| Pb | 65.0 | ug/L | o,p'-DDE | **<0.03** | ug/L |
| Cr | 55.6 | ug/L | p,p'-DDD | **<0.05** | ug/L |
| α-Endosulfan | **<0.03** | ug/L | p,p'-DDT | **<0.04** | ug/L |
| β-Endosulfan | **<0.04** | ug/L | | | |

**Table S3.**
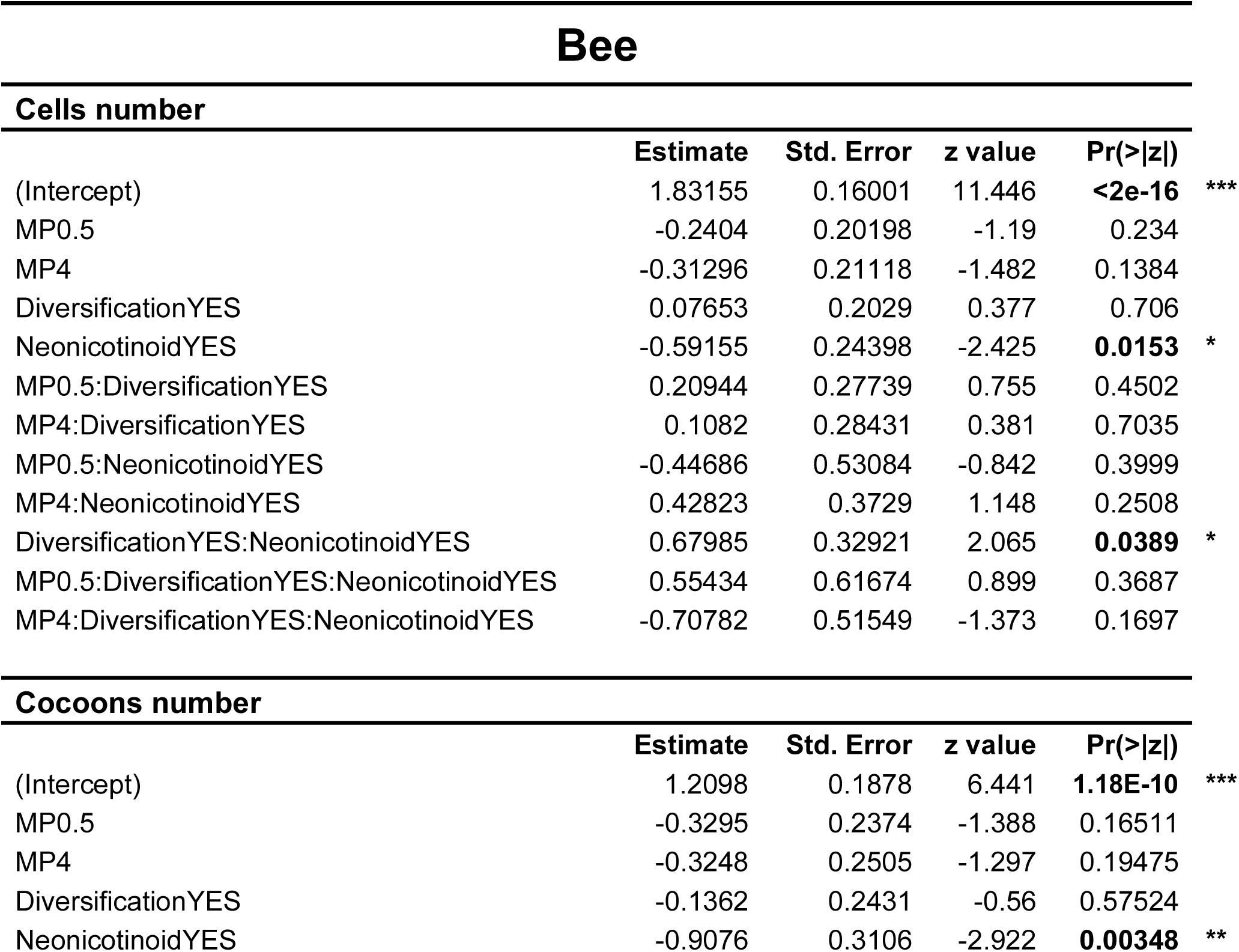
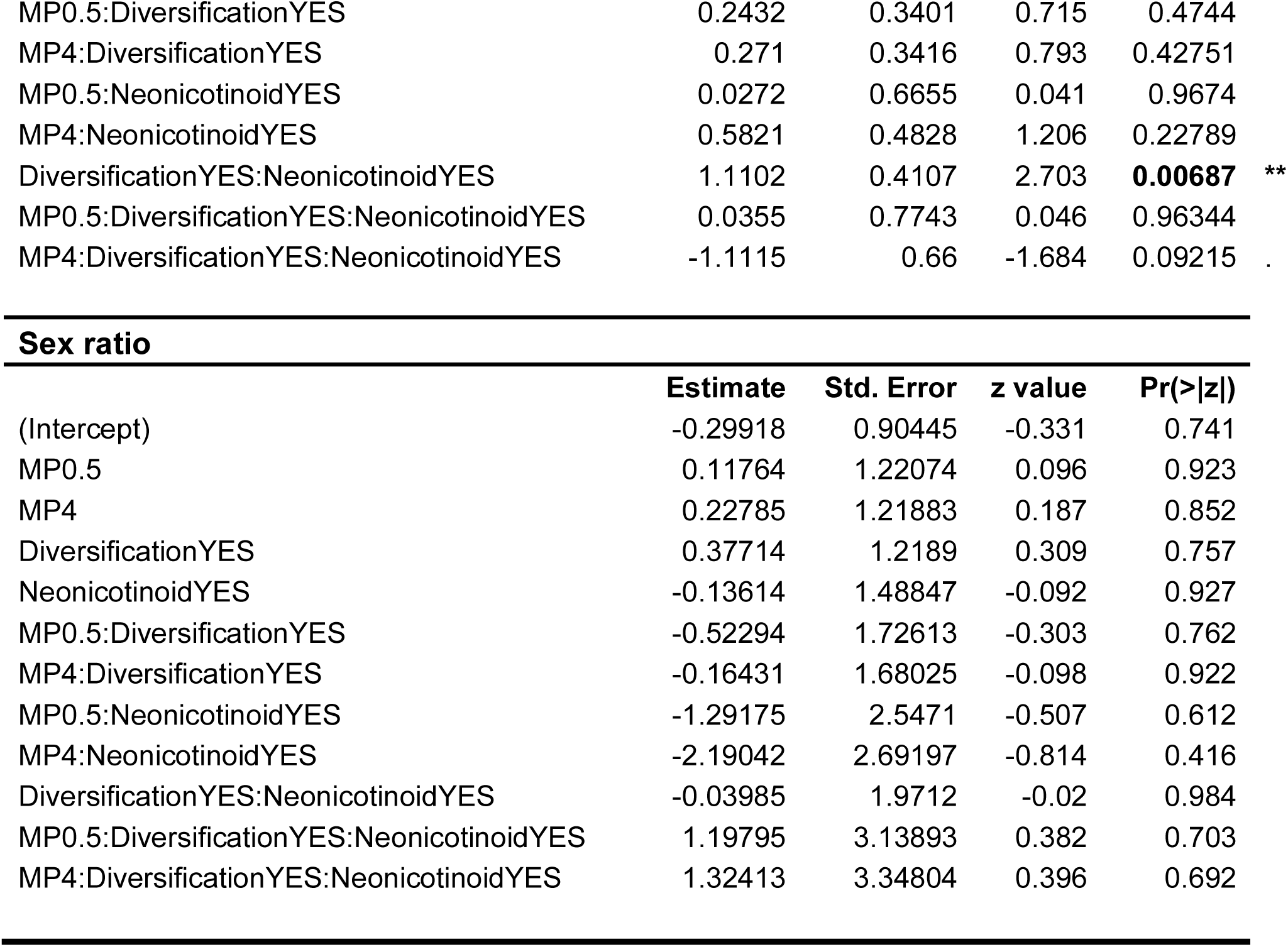
Detail of GLMM model results for bee trait in relation to treatment.

| **Bee** |  |  |  |  |  |
| --- | --- | --- | --- | --- | --- |
| **Cells number** |  |  |  |  |  |
|  | **Estimate** | **Std. Error** | **z value** | **Pr(>\|z\|)** |  |
| (Intercept) | 1.83155 | 0.16001 | 11.446 | **<2e-16** | *** |
| MP0.5 | -0.2404 | 0.20198 | -1.19 | 0.234 |  |
| MP4 | -0.31296 | 0.21118 | -1.482 | 0.1384 |  |
| DiversificationYES | 0.07653 | 0.2029 | 0.377 | 0.706 |  |
| NeonicotinoidYES | -0.59155 | 0.24398 | -2.425 | **0.0153** | * |
| MP0.5:DiversificationYES | 0.20944 | 0.27739 | 0.755 | 0.4502 |  |
| MP4:DiversificationYES | 0.1082 | 0.28431 | 0.381 | 0.7035 |  |
| MP0.5:NeonicotinoidYES | -0.44686 | 0.53084 | -0.842 | 0.3999 |  |
| MP4:NeonicotinoidYES | 0.42823 | 0.3729 | 1.148 | 0.2508 |  |
| DiversificationYES:NeonicotinoidYES | 0.67985 | 0.32921 | 2.065 | **0.0389** | * |
| MP0.5:DiversificationYES:NeonicotinoidYES | 0.55434 | 0.61674 | 0.899 | 0.3687 |  |
| MP4:DiversificationYES:NeonicotinoidYES | -0.70782 | 0.51549 | -1.373 | 0.1697 |  |
| **Cocoons number** |  |  |  |  |  |
|  | **Estimate** | **Std. Error** | **z value** | **Pr(>\|z\|)** |  |
| (Intercept) | 1.2098 | 0.1878 | 6.441 | **1.18E-10** | *** |
| MP0.5 | -0.3295 | 0.2374 | -1.388 | 0.16511 |  |
| MP4 | -0.3248 | 0.2505 | -1.297 | 0.19475 |  |
| DiversificationYES | -0.1362 | 0.2431 | -0.56 | 0.57524 |  |
| NeonicotinoidYES | -0.9076 | 0.3106 | -2.922 | **0.00348** | ** |
| MP0.5:DiversificationYES | 0.2432 | 0.3401 | 0.715 | 0.4744 |  |
| MP4:DiversificationYES | 0.271 | 0.3416 | 0.793 | 0.42751 |  |
| MP0.5:NeonicotinoidYES | 0.0272 | 0.6655 | 0.041 | 0.9674 |  |
| MP4:NeonicotinoidYES | 0.5821 | 0.4828 | 1.206 | 0.22789 |  |
| DiversificationYES:NeonicotinoidYES | 1.1102 | 0.4107 | 2.703 | **0.00687** | ** |
| MP0.5:DiversificationYES:NeonicotinoidYES | 0.0355 | 0.7743 | 0.046 | 0.96344 |  |
| MP4:DiversificationYES:NeonicotinoidYES | -1.1115 | 0.66 | -1.684 | 0.09215 | . |

Sex ratio
|  | Estimate | Std. Error | z value | Pr(>\|z\|) |
| --- | --- | --- | --- | --- |
| (Intercept) | -0.29918 | 0.90445 | -0.331 | 0.741 |
| MP0.5 | 0.11764 | 1.22074 | 0.096 | 0.923 |
| MP4 | 0.22785 | 1.21883 | 0.187 | 0.852 |
| DiversificationYES | 0.37714 | 1.2189 | 0.309 | 0.757 |
| NeonicotinoidYES | -0.13614 | 1.48847 | -0.092 | 0.927 |
| MP0.5:DiversificationYES | -0.52294 | 1.72613 | -0.303 | 0.762 |
| MP4:DiversificationYES | -0.16431 | 1.68025 | -0.098 | 0.922 |
| MP0.5:NeonicotinoidYES | -1.29175 | 2.5471 | -0.507 | 0.612 |
| MP4:NeonicotinoidYES | -2.19042 | 2.69197 | -0.814 | 0.416 |
| DiversificationYES:NeonicotinoidYES | -0.03985 | 1.9712 | -0.02 | 0.984 |
| MP0.5:DiversificationYES:NeonicotinoidYES | 1.19795 | 3.13893 | 0.382 | 0.703 |
| MP4:DiversificationYES:NeonicotinoidYES | 1.32413 | 3.34804 | 0.396 | 0.692 |

**Table S4.**
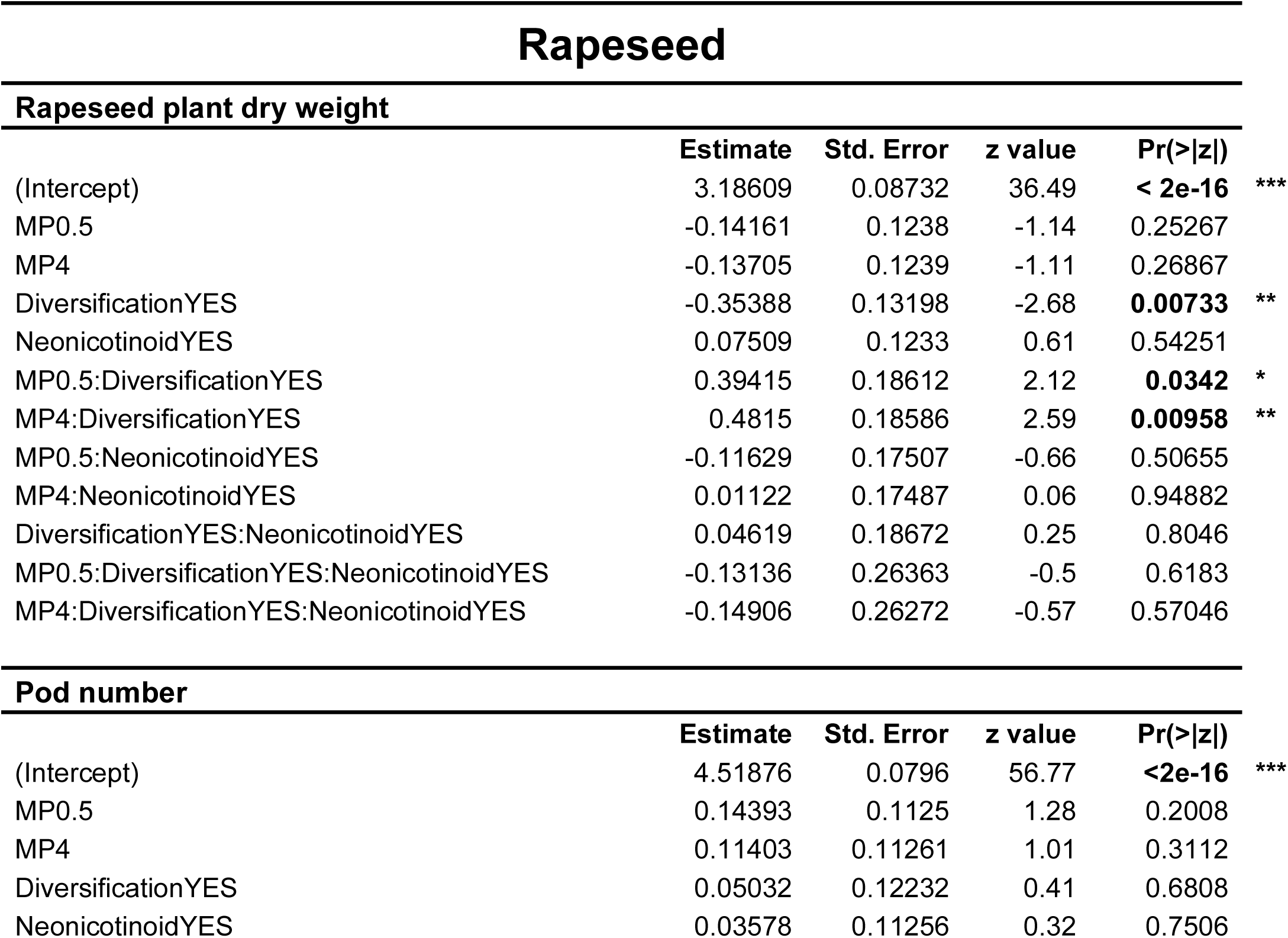
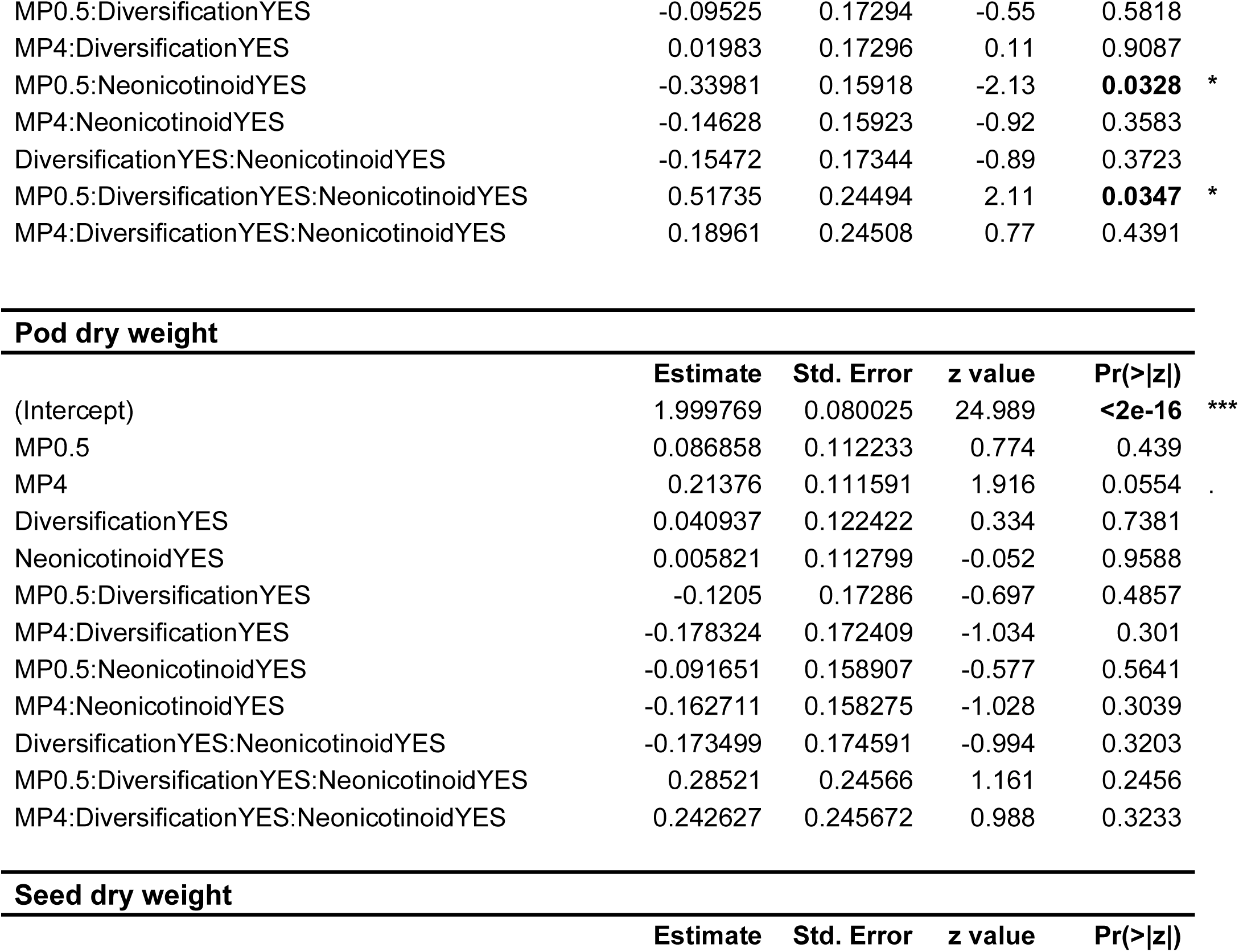
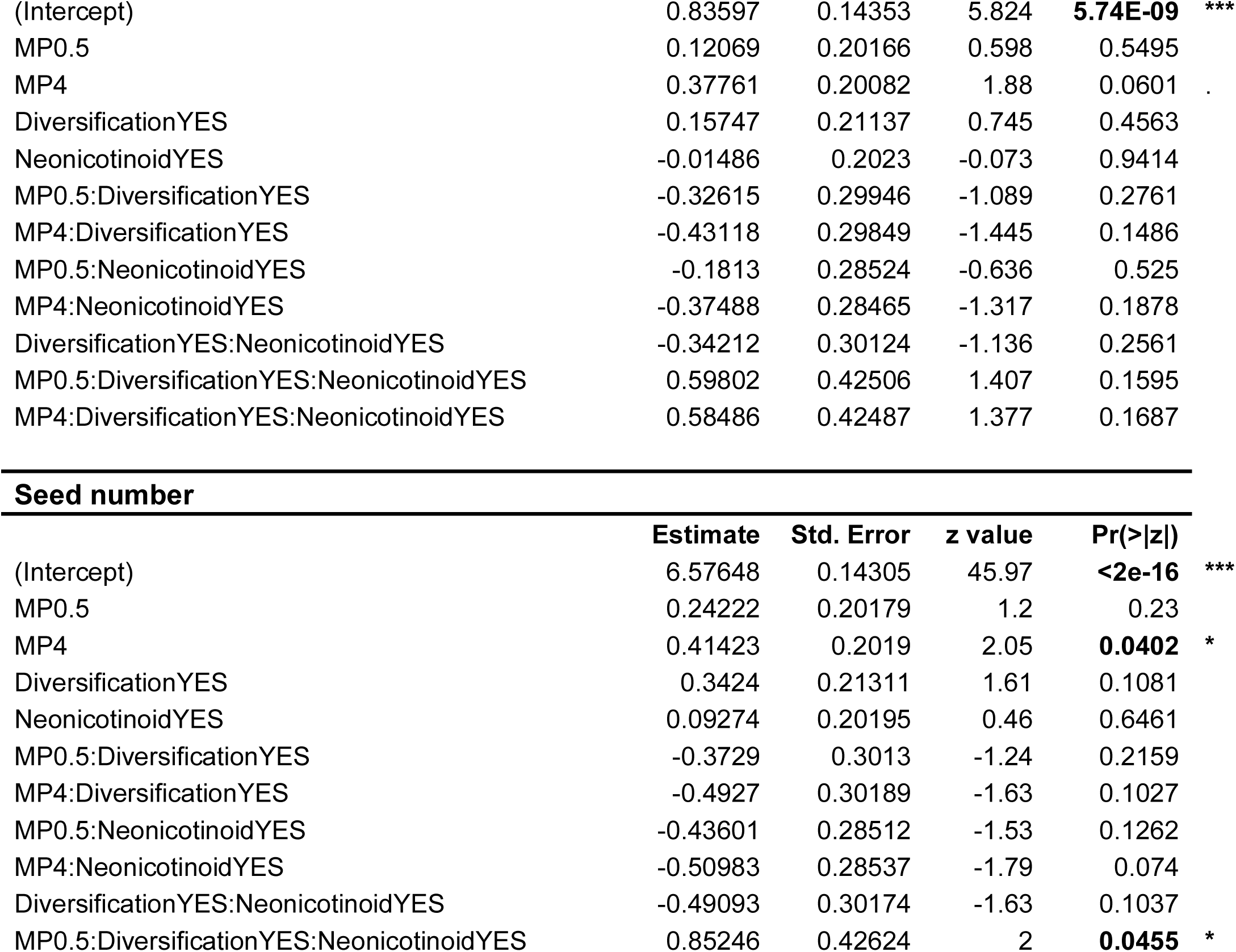
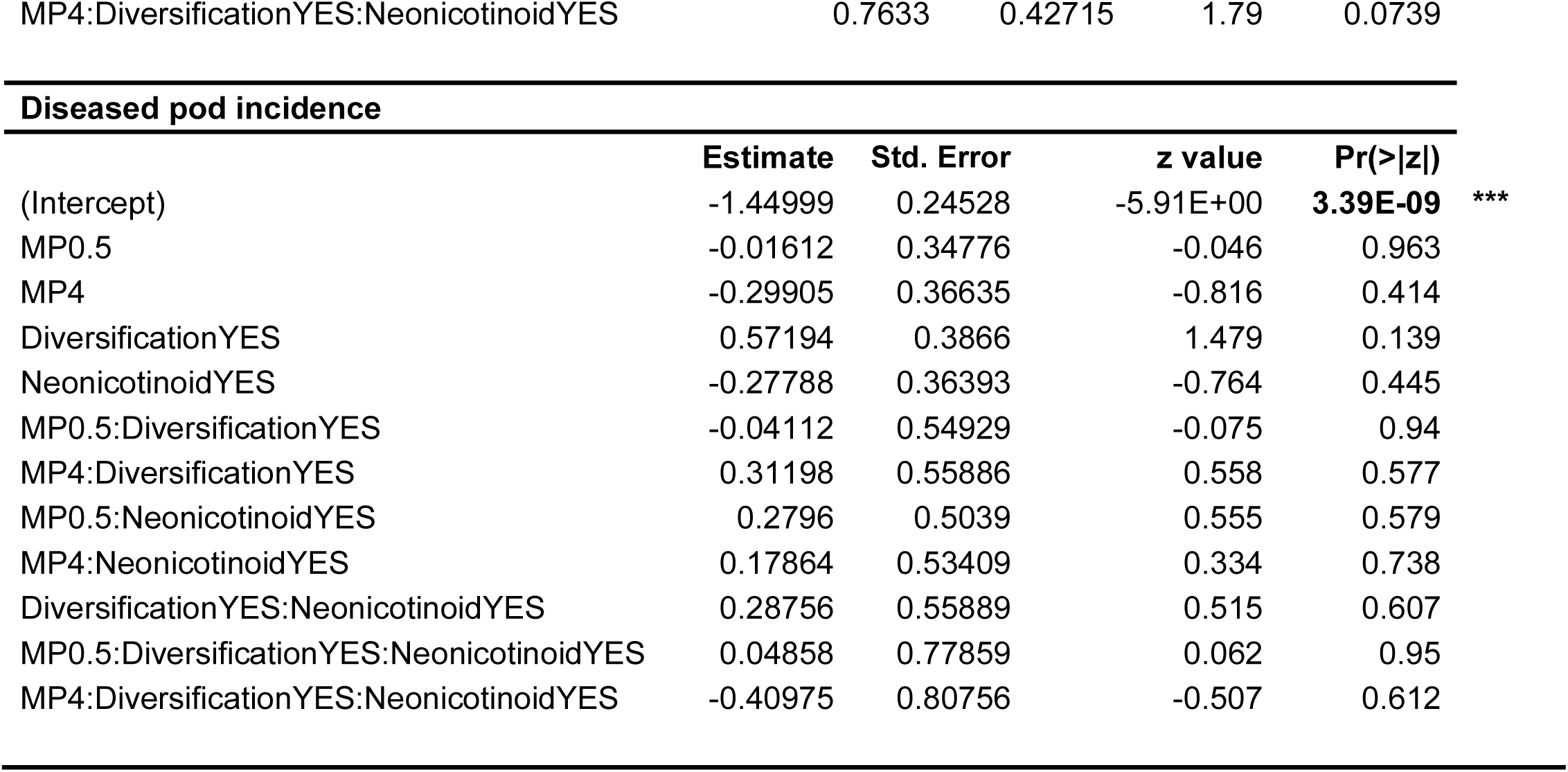
Detail of GLMM model results for rapeseed trait in relation to treatment.

**Table S5.** Detail of GAMM model results for rapeseed flower number in relation to treatment.

| **rapeseed flower number** | **Estimate** | **Std. Error** | **t value** | **Pr(>\|t\|)** |  |
| --- | --- | --- | --- | --- | --- |
| (Intercept) | 5.48055 | 0.13619 | 40.242 | **2.00E-16** | *** |
| NeonicotinoidYES | 0.04255 | 0.19259 | 0.221 | 0.825 |  |
| DiversificationYES | -0.99448 | 0.19389 | -5.129 | **4.47E-07** | *** |
| MP0.5 | -0.20594 | 0.19267 | -1.069 | 0.286 |  |
| MP4 | -0.20504 | 0.19266 | -1.064 | 0.288 |  |
| NeonicotinoidYES:DiversificationYES | 0.35529 | 0.274 | 1.297 | 0.195 |  |
| NeonicotinoidYES:MP0.5 | 0.17004 | 0.27242 | 0.624 | 0.533 |  |
| NeonicotinoidYES:MP4 | 0.13738 | 0.27242 | 0.504 | 0.614 |  |
| DiversificationYES:MP0.5 | 0.3306 | 0.27361 | 1.208 | 0.228 |  |
| DiversificationYES:MP4 | 0.39689 | 0.27412 | 1.448 | 0.148 |  |
| NeonicotinoidYES:DiversificationYES:MP0.5 | -0.44342 | 0.38672 | -1.147 | 0.252 |  |
| NeonicotinoidYES:DiversificationYES:MP4 | -0.45164 | 0.38709 | -1.167 | 0.244 |  |

**Table S6.**
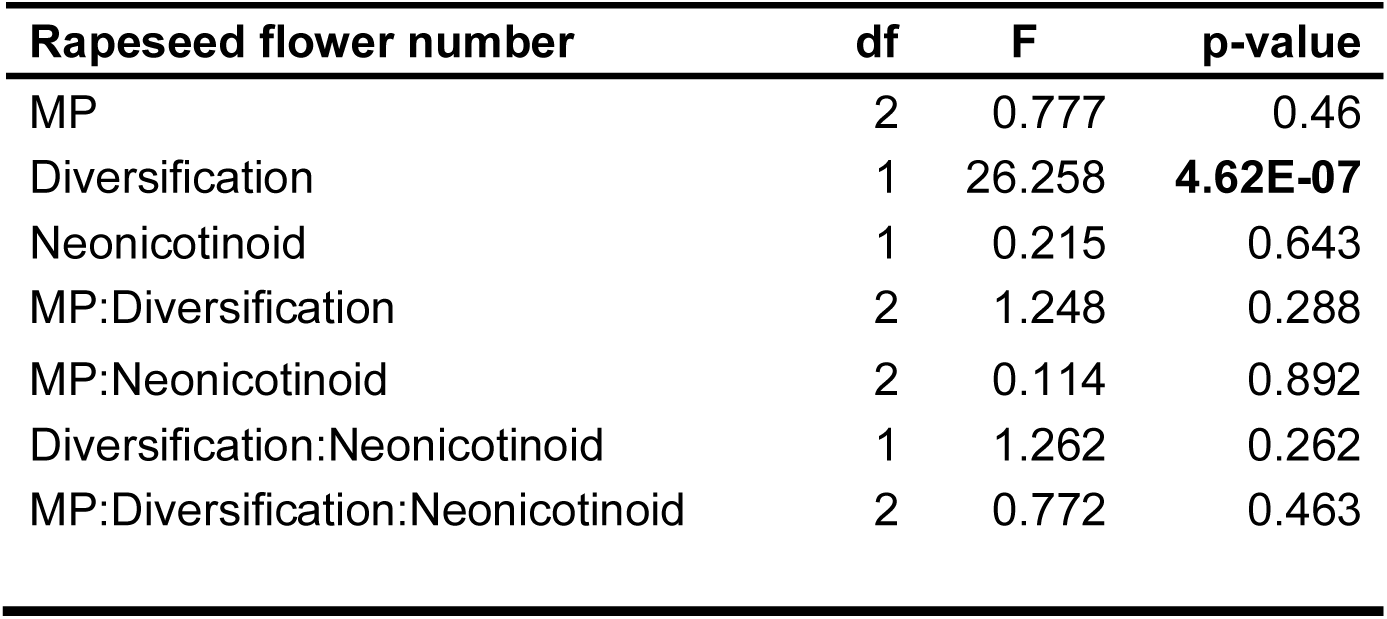
Anova test of testing the experimental treatment effects on rapeseed flower number.

| Rapeseed flower number | df | F | p-value |
| --- | --- | --- | --- |
| MP | 2 | 0.777 | 0.46 |
| Diversification | 1 | 26.258 | **4.62E-07** |
| Neonicotinoid | 1 | 0.215 | 0.643 |
| MP:Diversification | 2 | 1.248 | 0.288 |
| MP:Neonicotinoid | 2 | 0.114 | 0.892 |
| Diversification:Neonicotinoid | 1 | 1.262 | 0.262 |
| MP:Diversification:Neonicotinoid | 2 | 0.772 | 0.463 |

**Table S7.** Type-III Analysis of testing the experimental treatment effects on bee traits.

| **Bee** |  |  |  |
| --- | --- | --- | --- |
| **Cells number** |  |  |  |
|  | **Chisq** | **Df** | **Pr(>Chisq)** |
| MP | 3.6488 | 2 | 0.161314 |
| Diversification | 10.7358 | 1 | **0.001051** ** |
| Neonicotinoid | 2.053 | 1 | 0.151911 |
| MP:Diversification | 1.963 | 2 | 0.374742 |
| MP:Neonicotinoid | 0.0675 | 2 | 0.966802 |
| Diversification:Neonicotinoid | 5.8667 | 1 | **0.01543** * |
| MP:Diversification:Neonicotinoid | 4.0044 | 2 | 0.13504 |
| **Cocoons number** |  |  |  |
|  | **Chisq** | **Df** | **Pr(>Chisq)** |
| MP | 1.6678 | 2 | 0.434342 |
| Diversification | 3.1962 | 1 | 0.073811 |
| Neonicotinoid | 2.1968 | 1 | 0.1383 |
| MP:Diversification | 0.4548 | 2 | 0.796585 |
| MP:Neonicotinoid | 0.1916 | 2 | 0.908642 |
| Diversification:Neonicotinoid | 7.1109 | 1 | **0.007662** ** |
| MP:Diversification:Neonicotinoid | 3.2429 | 2 | 0.197608 |
| **Sex ratio** |  |  |  |
|  | **Chisq** | **Df** | **Pr(>Chisq)** |
| MP | 0.2772 | 2 | 0.8706 |
| Diversification | 0.3 | 1 | 0.5839 |
| Neonicotinoid | 0.9915 | 1 | 0.3194 |
| MP:Diversification | 0.0551 | 2 | 0.9728 |
| MP:Neonicotinoid | 0.7251 | 2 | 0.6959 |
| Diversification:Neonicotinoid | 0.2298 | 1 | 0.6317 |
| MP:Diversification:Neonicotinoid | 0.2206 | 2 | 0.8956 |

**Table S8.** Type-III Analysis of testing the experimental treatment effects on rapeseed traits.

| **Rapeseed** |  |  |  |
| --- | --- | --- | --- |
| **Rapeseed plant dry weight** |  |  |  |
|  | **Chisq** | **Df** | **Pr(>Chisq)** |
| MP | 2.5309 | 2 | 0.282106 |
| Diversification | 2.4738 | 1 | 0.115755 |
| Neonicotinoid | 0.124 | 1 | 0.724713 |
| MP:Diversification | 10.7694 | 2 | **0.004586** ** |
| MP:Neonicotinoid | 1.8507 | 2 | 0.39639 |
| Diversification:Neonicotinoid | 0.1978 | 1 | 0.656528 |
| MP:Diversification:Neonicotinoid | 0.3825 | 2 | 0.825918 |
| **Pod number** |  |  |  |
|  | **Chisq** | **Df** | **Pr(>Chisq)** |
| MP | 2.1683 | 2 | 0.3382 |
| Diversification | 1.7338 | 1 | 0.1879 |
| Neonicotinoid | 3.4645 | 1 | 0.0627 |
| MP:Diversification | 1.8588 | 2 | 0.3948 |
| MP:Neonicotinoid | 1.0077 | 2 | 0.6042 |
| Diversification:Neonicotinoid | 0.6625 | 1 | 0.4157 |
| MP:Diversification:Neonicotinoid | 4.5692 | 2 | 0.1018 |
| **Pod dry weight** |  |  |  |
|  | **Chisq** | **Df** | **Pr(>Chisq)** |
| MP | 3.2264 | 2 | 0.19925 |
| Diversification | 1.3086 | 1 | 0.25264 |
| Neonicotinoid | 3.3094 | 1 | 0.06888 |
| MP:Diversification | 0.4544 | 2 | 0.79676 |
| MP:Neonicotinoid | 0.5761 | 2 | 0.74971 |
| Diversification:Neonicotinoid | 0.0013 | 1 | 0.97103 |
| MP:Diversification:Neonicotinoid | 1.5641 | 2 | 0.45746 |

Seed dry weight
|  | Chisq | Df | Pr(>Chisq) |
| --- | --- | --- | --- |
| MP | 1.7171 | 2 | 0.42378 |
| Diversification | 0.6268 | 1 | 0.42852 |
| Neonicotinoid | 4.2327 | 1 | **0.03965** * |
| MP:Diversification | 0.5022 | 2 | 0.77795 |
| MP:Neonicotinoid | 0.9038 | 2 | 0.63641 |
| Diversification:Neonicotinoid | 0.0952 | 1 | 0.75771 |
| MP:Diversification:Neonicotinoid | 2.5792 | 2 | 0.27538 |

Seed number
|  | Chisq | Df | Pr(>Chisq) |
| --- | --- | --- | --- |
| MP | 1.0673 | 2 | 0.58647 |
| Diversification | 0.8059 | 1 | 0.36934 |
| Neonicotinoid | 5.4127 | 1 | **0.01999** * |
| MP:Diversification | 0.6189 | 2 | 0.73384 |
| MP:Neonicotinoid | 0.6561 | 2 | 0.72032 |
| Diversification:Neonicotinoid | 0.0754 | 1 | 0.78362 |
| MP:Diversification:Neonicotinoid | 4.8256 | 2 | 0.08956 |

**Diseased pod incidence**
|  | **Chisq** | **Df** | **Pr(>Chisq)** |  |
| --- | --- | --- | --- | --- |
| MP | 2.0262 | 2 | 0.3631 |  |
| Diversification | 21.1522 | 1 | **4.24E-06** | **\*\*\*** |
| Neonicotinoid | 0.0928 | 1 | 0.7606 |  |
| MP:Diversification | 0.1216 | 2 | 0.941 |  |
| MP:Neonicotinoid | 0.8079 | 2 | 0.6677 |  |
| Diversification:Neonicotinoid | 0.3045 | 1 | 0.5811 |  |
| MP:Diversification:Neonicotinoid | 0.3887 | 2 | 0.8234 |  |

**Table S9.** Detail of GAMM model results for all species flower number in relation to diversification treatment.

| **All flower number in all treatment** |  |  |  |  |  |
| --- | --- | --- | --- | --- | --- |
|  | **Estimate** | **Std. Error** | **t value** | **Pr(>\|t\|)** |  |
| (Intercept) | 3.06114 | 0.04307 | 71.08 | **<2e-16** | *** |
| DiversificationYES | -1.18239 | 0.05749 | -20.57 | **<2e-16** | *** |
| **All flower number in Control** |  |  |  |  |  |
|  | **Estimate** | **Std. Error** | **t value** | **Pr(>\|t\|)** |  |
| (Intercept) | 3.12018 | 0.07867 | 39.66 | **<2e-16** | *** |
| DiversificationYES | -1.42141 | 0.10364 | -13.71 | **<2e-16** | *** |

**Table S10.**
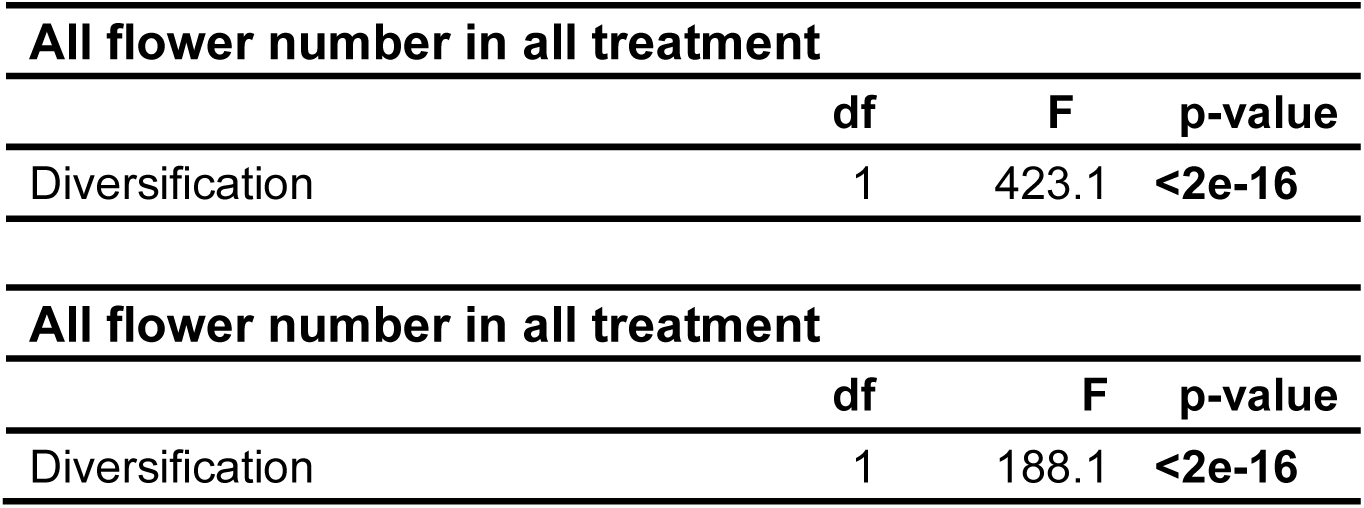
Anova test of testing the diversification treatment effects on all species flower number.

**Table S11.** Detail of GLMM model results for bee in relation to diversification. A total of three (species3) and four (species4) species of diversified flower resource were observed to bloom during the course of the experiment.

| Bee |  |  |  |  |  |  |  |  |  |  |  |
| --- | --- | --- | --- | --- | --- | --- | --- | --- | --- | --- | --- |
| Shannon Index |  |  |  |  |  | Species Number |  |  |  |  |  |
| Response variable |  |  |  |  |  |  |  |  |  |  |  |
| Cells number<br>(n=206) |  |  |  |  |  | (n=206) |  |  |  |  |  |
|  | Estimate | Std. Error | z value | Pr(>\|z\|) |  |  | Estimate | Std. Error | z value | Pr(>\|z\|) |  |
| (Intercept) | 1.4712 | 0.07818 | 18.817 | < 2e-16 | *** | (Intercept) | 1.47219 | 0.07996 | 18.412 | < 2e-16 | *** |
| shannon | **0.61144** | **0.16518** | **3.702** | **0.000214** | *** | species3 | 0.24724 | 0.30575 | 0.809 | 0.418724 |  |
|  |  |  |  |  |  | species4 | **0.38792** | **0.10925** | **3.551** | **0.000384** | *** |
| Cocoons number |  |  |  |  |  |  |  |  |  |  |  |
| (n=206) |  |  |  |  |  | (n=206) |  |  |  |  |  |
|  | Estimate | Std. Error | value | Pr(>\|z\|) |  |  | Estimate | Std. Error | z value | Pr(>\|z\|) |  |
| (Intercept) | 0.79227 | 0.08647 | 9.162 | < 2e-16 | *** | (Intercept) | 0.80987 | 0.08702 | 9.307 | < 2e-16 | *** |
| shannon | **0.50765** | **0.18048** | **2.813** | **0.00491** | ** | species3 | -0.32983 | 0.45399 | -0.727 | 0.46753 |  |
|  |  |  |  |  |  | species4 | **0.30712** | **0.11673** | **2.631** | **0.00852** | ** |

**Table S12.**
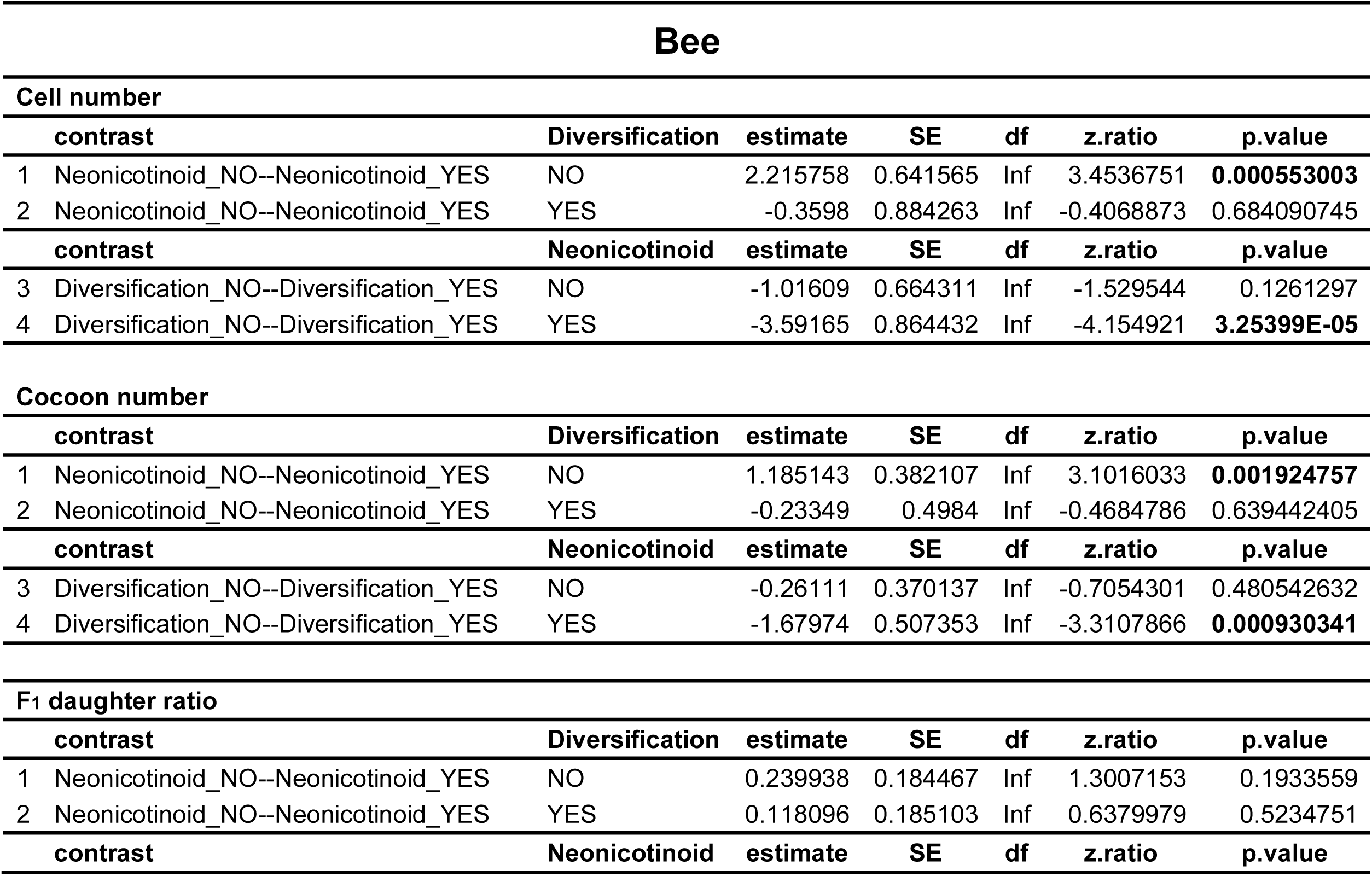
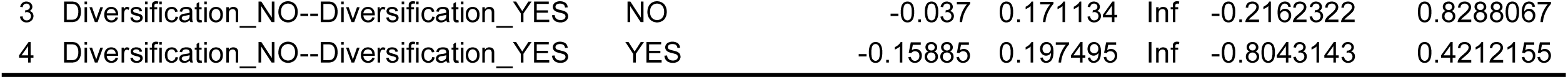
Detail of pairwise comparisons using Tukey’s HSD adjustment for bee traits in relation to neonicotinoid and diversification.

| **Bee** |  |  |  |  |  |  |
| --- | --- | --- | --- | --- | --- | --- |
| **Cell number** |  |  |  |  |  |  |
| **contrast** | **Diversification** | **estimate** | **SE** | **df** | **z.ratio** | **p.value** |
| 1 Neonicotinoid_NO--Neonicotinoid_YES | NO | 2.215758 | 0.641565 | Inf | 3.4536751 | **0.000553003** |
| 2 Neonicotinoid_NO--Neonicotinoid_YES | YES | -0.3598 | 0.884263 | Inf | -0.4068873 | 0.684090745 |
| **contrast** | **Neonicotinoid** | **estimate** | **SE** | **df** | **z.ratio** | **p.value** |
| 3 Diversification_NO--Diversification_YES | NO | -1.01609 | 0.664311 | Inf | -1.529544 | 0.1261297 |
| 4 Diversification_NO--Diversification_YES | YES | -3.59165 | 0.864432 | Inf | -4.154921 | **3.25399E-05** |
| **Cocoon number** |  |  |  |  |  |  |
| **contrast** | **Diversification** | **estimate** | **SE** | **df** | **z.ratio** | **p.value** |
| 1 Neonicotinoid_NO--Neonicotinoid_YES | NO | 1.185143 | 0.382107 | Inf | 3.1016033 | **0.001924757** |
| 2 Neonicotinoid_NO--Neonicotinoid_YES | YES | -0.23349 | 0.4984 | Inf | -0.4684786 | 0.639442405 |
| **contrast** | **Neonicotinoid** | **estimate** | **SE** | **df** | **z.ratio** | **p.value** |
| 3 Diversification_NO--Diversification_YES | NO | -0.26111 | 0.370137 | Inf | -0.7054301 | 0.480542632 |
| 4 Diversification_NO--Diversification_YES | YES | -1.67974 | 0.507353 | Inf | -3.3107866 | **0.000930341** |
| **F1 daughter ratio** |  |  |  |  |  |  |
| **contrast** | **Diversification** | **estimate** | **SE** | **df** | **z.ratio** | **p.value** |
| 1 Neonicotinoid_NO--Neonicotinoid_YES | NO | 0.239938 | 0.184467 | Inf | 1.3007153 | 0.1933559 |
| 2 Neonicotinoid_NO--Neonicotinoid_YES | YES | 0.118096 | 0.185103 | Inf | 0.6379979 | 0.5234751 |
| **contrast** | **Neonicotinoid** | **estimate** | **SE** | **df** | **z.ratio** | **p.value** |

|  |  |  |  |  |  |  |  |
| --- | --- | --- | --- | --- | --- | --- | --- |
| 3 | Diversification_NO--Diversification_YES | NO | -0.037 | 0.171134 | Inf | -0.2162322 | 0.8288067 |
| 4 | Diversification_NO--Diversification_YES | YES | -0.15885 | 0.197495 | Inf | -0.8043143 | 0.4212155 |

**Table S13.**
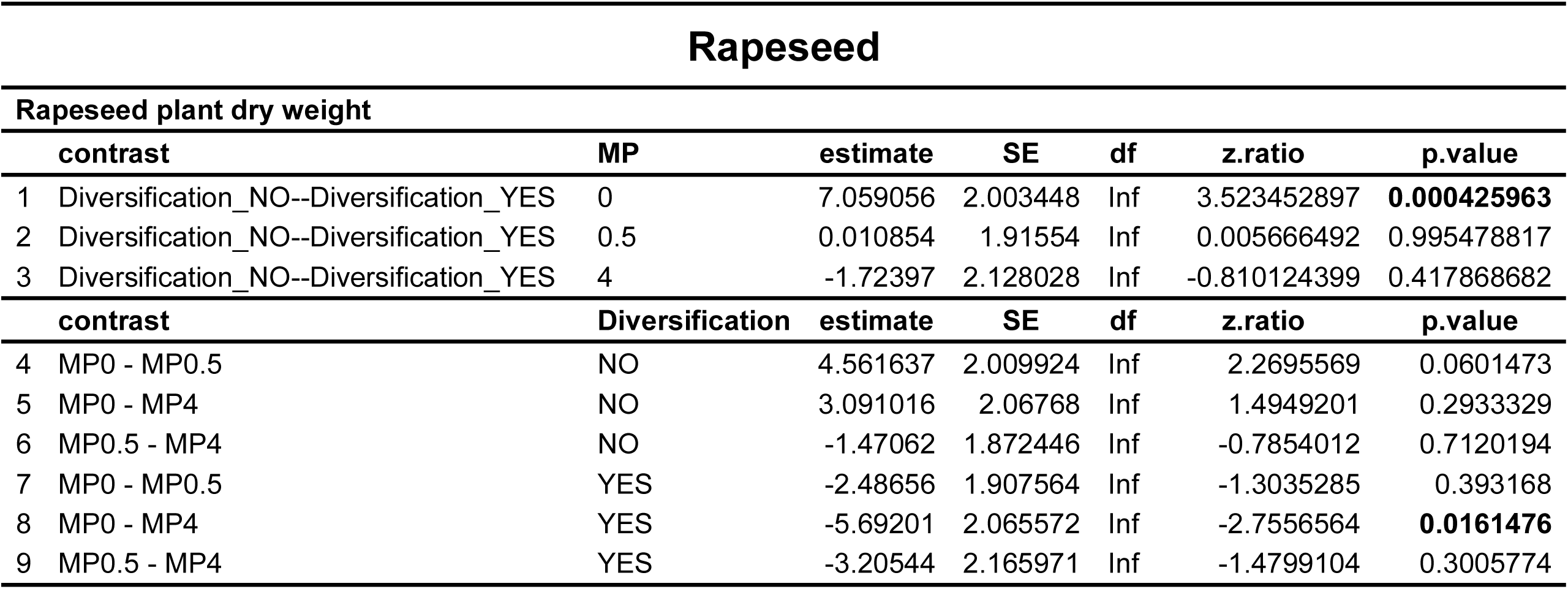
Detail of pairwise comparisons using Tukey’s HSD adjustment for rapeseed dry weight in relation to microplastic and diversification.

| Rapeseed |  |  |  |  |  |  |
| --- | --- | --- | --- | --- | --- | --- |
| Rapeseed plant dry weight |  |  |  |  |  |  |
| contrast | MP | estimate | SE | df | z.ratio | p.value |
| 1 Diversification_NO--Diversification_YES | 0 | 7.059056 | 2.003448 | Inf | 3.523452897 | **0.000425963** |
| 2 Diversification_NO--Diversification_YES | 0.5 | 0.010854 | 1.91554 | Inf | 0.005666492 | 0.995478817 |
| 3 Diversification_NO--Diversification_YES | 4 | -1.72397 | 2.128028 | Inf | -0.810124399 | 0.417868682 |
| contrast | Diversification | estimate | SE | df | z.ratio | p.value |
| 4 MP0 - MP0.5 | NO | 4.561637 | 2.009924 | Inf | 2.2695569 | 0.0601473 |
| 5 MP0 - MP4 | NO | 3.091016 | 2.06768 | Inf | 1.4949201 | 0.2933329 |
| 6 MP0.5 - MP4 | NO | -1.47062 | 1.872446 | Inf | -0.7854012 | 0.7120194 |
| 7 MP0 - MP0.5 | YES | -2.48656 | 1.907564 | Inf | -1.3035285 | 0.393168 |
| 8 MP0 - MP4 | YES | -5.69201 | 2.065572 | Inf | -2.7556564 | **0.0161476** |
| 9 MP0.5 - MP4 | YES | -3.20544 | 2.165971 | Inf | -1.4799104 | 0.3005774 |

